# Signalling involving MET and FAK supports cell division independent of the activity of the cell cycle-regulating CDK4/6 kinases

**DOI:** 10.1101/601799

**Authors:** Chi Zhang, Simon R. Stockwell, May Elbanna, Robin Ketteler, Jamie Freeman, Bissan Al-Lazikani, Suzanne Eccles, Alexis De Haven Brandon, Florence Raynaud, Angela Hayes, Paul A. Clarke, Paul Workman, Sibylle Mittnacht

**Affiliations:** UCL Cancer Institute, University College London, London, WC1E 6DD, UK; Cancer Research UK Cancer Therapeutics Unit at The Institute of Cancer Research, London, SM2 5NG; MRC Laboratory for Molecular Cell Biology, University College London, London, WC1E 6BT, UK

**Author notes:** **corresponding authors** Paul Workman, Cancer Research UK Cancer Therapeutics Unit at The Institute of Cancer Research, London, SM2 5NG., AND Sibylle Mittnacht, UCL Cancer Institute, University College London, London, WC1E 6DD, UK.

**Keywords:** CDK4, CDK6, CDK4/6 inhibitor, therapy-resistance, HGF, MET, FAK, combination therapy

## Abstract

Deregulation of the cyclin-dependent kinases 4 and 6 (CDK4/6) is highly prevalent in cancer yet inhibitors against these kinases currently show use in restricted tumour contexts. The extent to which cancers depend on CDK4/6 and what may undermine such dependency is poorly understood. Here we report that signalling engaging the MET proto-oncogene receptor tyrosine kinase/focal adhesion kinase (FAK) axis leads to CDK4/6-independent CDK2-activation, involving as a critical mechanistic events loss of the CDK inhibitor p21^CIP1^ and gain of its regulator, the ubiquitin ligase subunit SKP2. Combined inhibition of MET/FAK and CDK4/6 eliminates proliferation capacity of cancer cells in culture, and enhances tumour growth inhibition *in vivo*. Activation of the MET/FAK axis is known to arise through cancer extrinsic and intrinsic cues. Our work predicts that such cues support cell division independent of the activity of the cell cycle-regulating CDK4/6 kinases and identifies MET/FAK as a tractable route to broaden the utility of CDK4/6 inhibitor-based therapies in the clinic.

## Introduction

The cyclin-dependent kinases CDK4 and CDK6 are core components of the signal transduction network controlling transition of cells from G1 (Gap1) phase of the cell cycle into S (DNA synthesis) [1, 2]. Deregulation of this is a common event in cancer. Multiple oncogenic pathways promote the synthesis of the activator D-type cyclins, and gene mutation of regulators involved in limiting CDK4/6-activation are exceptionally frequent in all types of cancer [3, 4]. The high frequency by which CDK4/6 control is compromised in cancer implies that CDK4/6 deregulation is a key event enabling cancer development and, by extension, that inhibition of CDK4/6 could be a broadly applicable and effective approach to cancer treatment [2, 5].

Several potent, selective inhibitors targeting CDK4/6 (CDK4/6is) have undergone clinical trials including palbociclib (PD0332991), abemaciclib (LY-2835219) and ribociclib (LEE001) [6, 7], and have gained regulatory approval in combination with hormonal therapy in breast cancer [8–11]. However, evidence for clinical benefit has not been extended to other cancer types thus far, and relapse under therapy is frequent in the approved indication in breast cancer [9].

Activation of CDK4/6 requires their binding to D-type cyclins, synthesised in response to mitogenic signals [12, 13]. Bound to these cyclins CDK4/6 phosphorylates the retinoblastoma tumour suppressor protein (RB1), initiating its inactivation. RB1 in its active form prevents the transcription of genes required for S phase entry including those encoding the E-and A-type cyclins, involved in the activation of the CDK4/6 related S phase cyclin-dependent kinase CDK2 [14]. In addition RB1 promotes the ubiquitin-dependent destruction of the SCF (SKP1–CUL1–F-box protein) E3-ubiquitin-ligase substrate-recognition subunit SKP2, stabilizing the CDK2 inhibitory proteins p21^CIP1^ and p27^KIP1^, the degradation of which is SKP2-dependent [15, 16]. Together these activities limit CDK2-activation safeguarding licensed DNA synthesis and cell cycle transit.

Numerous reports describe situations where activation of CDK2 is enabled in the absence of CDK4/6 activity and that CDK2 can drive cycle transit in the absence CDK4/6 activity [17–24]. Hence activation of CDK2, independent of CDK4/6 activity, may limit the potency of CDK4/6is in cancers and identification of signalling required for CDK2-activation may yield information that predicts, or could be exploited to extend their efficacy in cancer therapy.

Here we report a mechanism-focussed screen aimed at identifying signalling that enables CDK4/6-independent CDK2-activation. We identify a prominent role of the MET proto-oncogene tyrosine kinase receptor family and their downstream effectors, the focal adhesion kinase (FAK) family. Our data validate MET/FAK signalling as a mechanism that enables CDK4/6-independent CDK2-activation and cell cycle transit and we provide evidence for the utility of MET-inhibition as a means to improve tumour response to CDK4/6-inhibition in vivo.

## Results

### Screening identifies proteins required for CDK2-activation in CDK4/6-inhibited cells

To assess CDK2-activation in cells exposed CDK4/6i), we used a cell-based CDK2 reporter (GFP-PSLD) where a green fluorescent protein (GFP) is fused to the CDK2-regulated phosphorylated subcellular localization domain (PSLD) of human DNA helicase B [25]. Phosphorylation of the PSLD by CDK2 exposes a nuclear export sequence initiating export of the GFP fusion protein from the nucleus to the cytoplasm (Fig. 1a). We made use of human colorectal carcinoma HCT116 cells (HCT116) that stably express the CDK2 reporter (HCT116-PSLD) [26]. Treatment of HCT116-PSLD using either siRNA targeting CDK2 or the CDK4/6i palbociclib significantly increased the fraction of cells with predominantly nuclear fluorescence (nuclear: cytoplasmic (nuc/cyto) fluorescence ratio > 1.5), consistent with reliance of reporter localization on CDK2, and CDK4/6-activation (Supplementary Fig. 1a-c). However, although palbociclib treatment increased the percentage of cells with nuclear-localised CDK2 reporter, a considerable portion of cells with loss of CDK4/6 activity, detected by the absence of RB1 phosphorylated at the CDK4/6-selective phosphorylation site Ser780 (pRB1^S780^) [27], continued to contain reporter with predominant cytoplasmic localization (Supplementary Fig. 1d, e). This indicates CDK4/6-independent CDK2-activation. Unexpectedly we observed that ablation of the tumour suppressor *TP53* reduced CDK2 control by palbociclib, even though the cells remained responsive to CDK4/6- inhibition, indicated by the reduction in cells containing pRB1^S780^ (Supplementary Fig. S1f and S1g).

**Figure 1.**
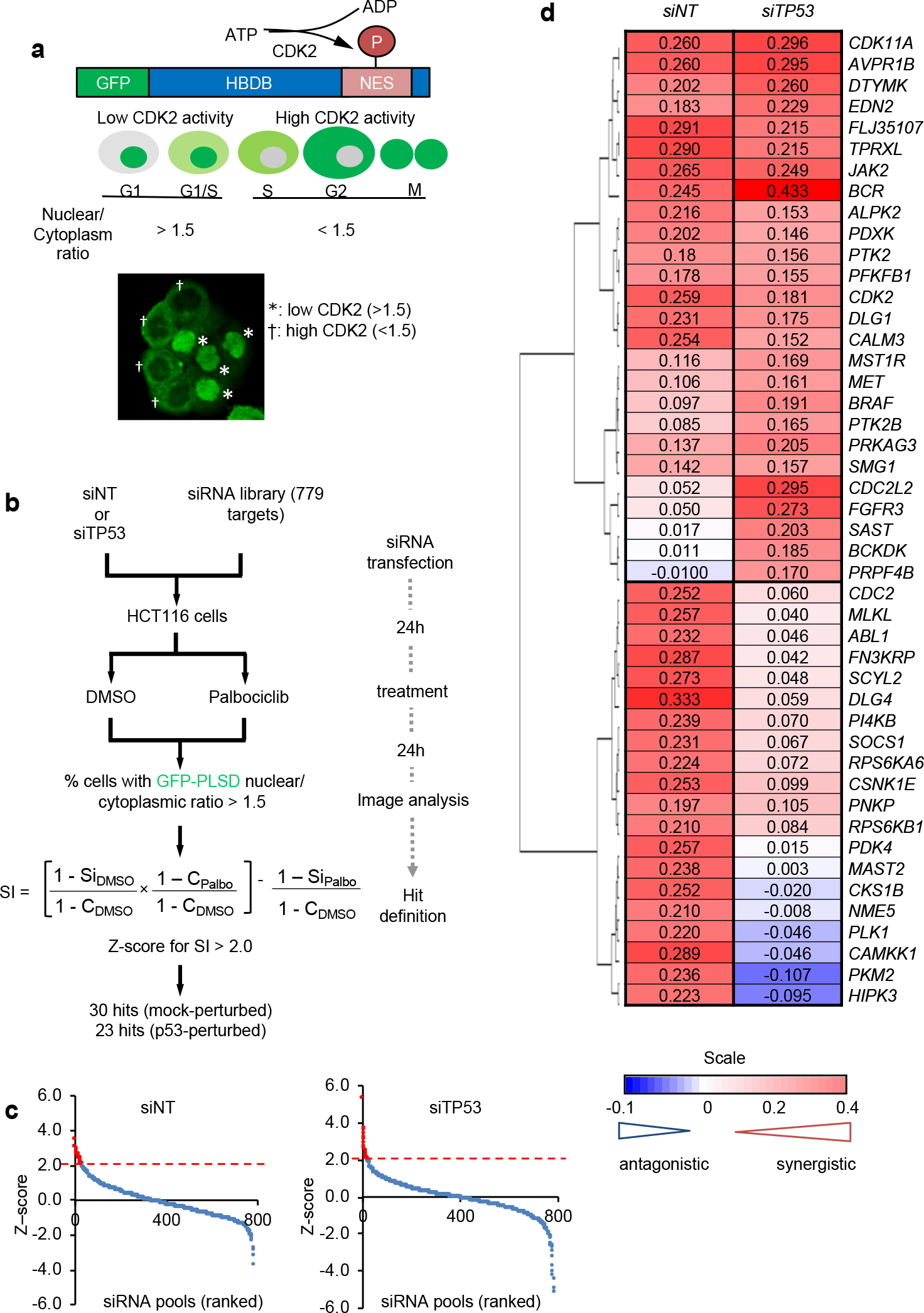
Screen for proteins permitting CDK2-activation in cells with CDK4/6- inhibition. **a** Schematic depicting functioning of the CDK2 reporter GFP-PSLD. Modular reporter structure, relationship between subcellular distribution of GFP and cell cycle phase, and a representative image of individual HCT116-PSLD with low (GFP-PSLD nuc/cyto > 1.5) or high (GFP-PSLD nuc/cyto < 1.5) CDK2 activity is shown. HDHB, human DNA helicase B; NES, nuclear export sequence. **b** Screen outline and procedure for hit identification. **c** Z-score ranking for siRNA pools in the screen. Results for unperturbed (siNT) and TP53-perturbed (siTP53) conditions are shown. Data points represent the mean of n=3 independent repeats, siRNA pools with Z-score > 2 marked in red. **d** Hierarchical clustering of hits based on mean (n = 3) sensitivity index values (SI). Colours denote the nature of interaction between siRNA pool and palbociclib: red, synergistic; white, additive; blue, antagonistic. siRNA target genes on the left. (Related to Supplementary Fig. 1)

To identify signalling that permits CDK4/6-independent CDK2 activation, we transfected HCT116-PSLD with a library of small interfering RNA (siRNA) pools targeting kinases and kinase-relevant components (Fig. 1B) and assessed the ability of palbociclib to restrain CDK2-activation under these conditions (Fig. 1b). Since we observed relaxed CDK2 control following ablation of *TP53* and because functional TP53 loss is frequent in cancer, we included an arm to the screen where we compromised TP53 expression using *TP53*-targeted siRNA.

To identify siRNA pools that selectively decreased CDK2 activity subject to palbociclib treatment we computed the sensitivity index (SI), which quantifies the difference between expected combined and measured observed effects of two treatments [28, 29], in this case the effects of specific siRNAs and the effect of CDK4/6-inhibition on CDK2 activity. Ranking the SI values calculated using Z-score statistics (Fig. 1c) we selected siRNA pools with Z-scores > 2 for further analysis, yielding 30 pools that selectively decreased CDK2 in combination with CDK4/6-inhibition in HCT116-PSLD, and 23 in HCT116-PSLD with compromised *TP53*-expression.

Most siRNA pools identified in *TP53*-compromised HCT116-PSLD decreased CDK2 activity also in HCT116-PSLD. Conversely, less than half identified in HCT116-PSLD decreased CDK2 activity in *TP53*-modified cells (Fig. 1d). These results indicate differences in the regulation of CDK2 in *TP53*-normal and *TP53*-imparied backgrounds but at the same time outline opportunity to enhance the dependence of CDK2-activation on CDK4/6 in cells, regardless of *TP53* status.

### MET/FAK signalling is required for CDK2-activation in CDK4/6-inhibited cells

To mine for annotated pathways overrepresented amongst the siRNA targets identified against those screened we used the MetaCore™ GeneGO tool (Supplementary tables 1 and 2), predicting as most significantly enriched a well-connected hub involving the MET proto-oncogene/hepatocyte growth factor receptor (MET) and the closely related macrophage growth factor receptor (MSTR1/RON) along with fibroblast growth factor receptor 3 (FGFR3) and their common downstream signalling targets, the focal adhesion kinases PTK2 and PTK2B (Fig. 2a).

**Figure 2.**
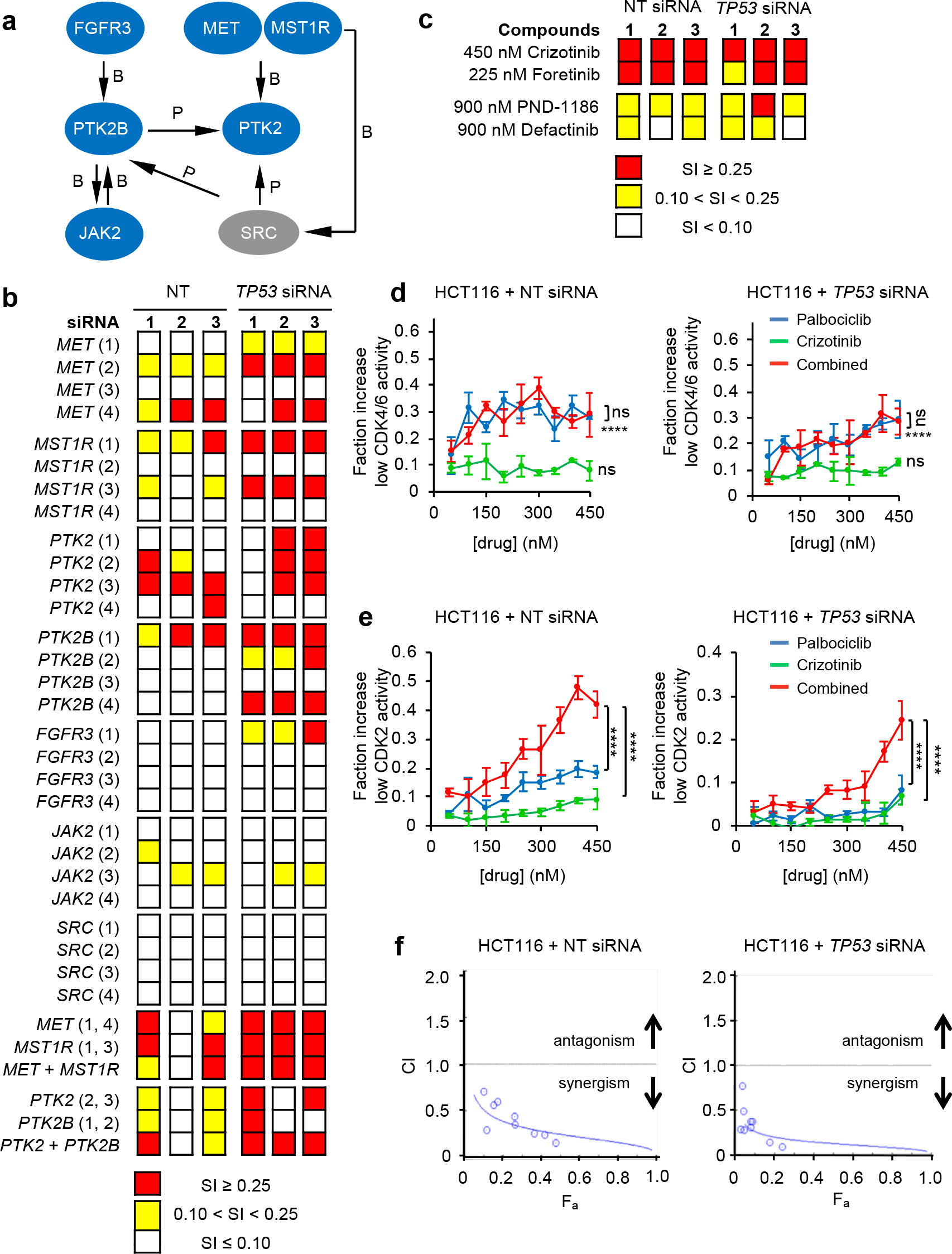
Signalling involving MET permits CDK2-activation in cells with CDK4/6- inhibition. **a** MetaCore™ GeneGO analysis identifies signalling network engaging MET overrepresented by hits. Interaction types: P, phosphorylation; B, binding; proteins targeted by screen identified siRNA pools in blue. **b-c** Hit validation using individual siRNAs (**b**) or pharmacological inhibitors for MET/MST1R (crizotinib or foretinib) or PTK2/2B (PDN-1186 or defactinib) (**c**). Data depict SI score relating to loss of CDK2 activity in combination with CDK4/6-inhibiton using palbociclib, determined using GFP-PSLD localisation. Data are mean ±SD for n=3 independent repeats. **d-e** Concentration-effect analysis depicting change in the fraction of cells with low CDK4/6 activity, assessed using automated microscopy analysis of cells immunostained for pRB1^S780^ (**d**) or low CDK2 activity, assessed based on GFP-PSLD localisation (**e**)following individual or combined inhibition of CDK4/6, using palboclib, and MET, using crizotinib. Data are mean ±SD for independent repeats. \*\*\*\**p*≤ 0.0001, ^ns^*p*> 0.05, 1-way ANOVA comparing effects across concentrations against vehicle or comparing palbociclib alone against palbociclib plus crizotinib (**d**) and comparing palbociclib alone or crizotinib alone against palbociclib plus crizotinib (**e**). **f** CI value plot calculated from data in E. F_a_= fraction affected.

Two or more distinct siRNAs targeting MET, MST1R, PTK2 or PTK2B synergistically increased the percentage cells with nuclear-localised PSLD-GFP following CDK4/6- inhibition (Fig. 2b) validating the involvement MET and FAK family members in enabling CDK4/6-independent CDK2-activation in cells. Combined use of *MET* and *MST1R* or *PTK2* and *PTK2B* siRNA did not enhance outcome, suggesting an independent, rate-limiting contribution of individual MET and FAK family kinases in this context. Notably, treatment with chemical inhibitors targeting either the MET or FAK family kinases synergistically decreased CDK2 activity in combination with palbociclib (Fig. 2c). The activity of network components FGFR3, SRC and JAK did not confirm with multiple oligonucleotides (Fig. 2b). Hence the involvement of these components in enabling CDK4/6-independent CDK2-activation cannot be certain.

To assess if inhibition of MET enables CDK2 control by enhancing the efficacy of the inhibitor to control CDK4/6, we assessed loss of pRB1^S780^ (Fig. 2d) in HCT116-PSLD subject to individual and combined use of inhibitors. As expected we observed a significant increase in the fraction of pRB1^S780^ negative cells following CDK4/6-inhibition. Conversely, MET-inhibition did not significantly increase the fraction of pRB1^S780^ negative cells. Importantly, combined inhibition of CDK4/6 and MET was no more effective at raising the fraction of pRB1^S780^ negative cells than inhibition of CDK4/6 alone at any concentration tested. Nevertheless, and in agreement with our earlier results, combined inhibition of CDK4/6 and MET led to a significantly greater reduction of cells with active CDK2 than treatment with either inhibitor alone (Fig. 2e). Chou-Talalay concentration-effect analysis [30] identified a robust synergistic interaction between MET and CDK4/6-inhibition towards reducing CDK2 activity, returning combination index (CI) values well below 1 across the concentration range tested (Fig. 2e)—irrespective of *TP53* status. Hence, MET-inhibition cooperates with palbociclib to control CDK2-activation but does not itself affect, or enhance, the ability of CDK4/6is to supress CDK4/6 activity.

### Combined MET and CDK4/6-inhibition synergistically affects tumour cell fate *in vitro* and reduces tumour growth *in vivo*

Since MET-inhibition synergised with CDK4/6-inhibition to enable the control of CDK2 activity, we tested if this treatment would also synergise to enable other responses associated with CDK4/6-inhibition. Inhibition of CDK4/6 is recognised for its ability to trigger permanent cell cycle exit, thought to underlie its anticancer activity [31]. To assess if MET inhibition enhances permanent cell cycle exit subject to CDK4/6-inhibition, we exposed cells for five consecutive days to inhibitors, then quantified their ability to form colonies by seeding equal numbers of live cells into inhibitor-free medium (Fig. 3). We initially measured the response of HCT116, using two chemically unrelated inhibitors for CDK4/6, palbociclib and abemaciclib, and chemically unrelated inhibitors of MET family kinases, crizotinib and foretinib (Fig. 3a-d), in accordance with best practice [32].

**Figure 3.**
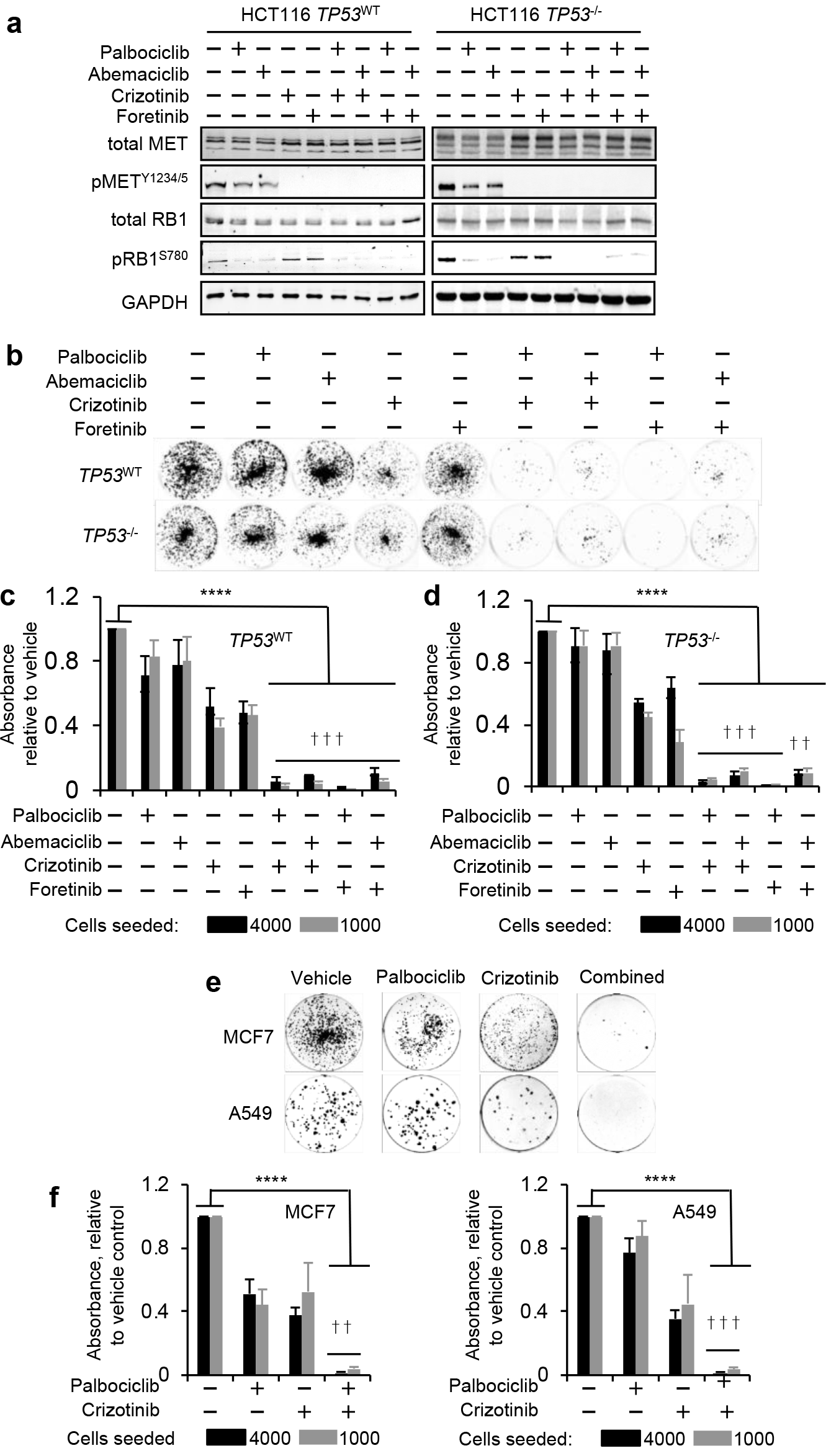
CDK4/6 and METis cooperate to incapacitate cell reproduction. **a-f** Clonogenic activity in cells with inhibitors for CDK4/6 (palbociclib or abermaciclib) or MET (crizotinib or foretinib). Clonogenic activity determined 120 h following inhibitor exposure. Data for HCT116 TP53^WT^ and TP53^−/−^ (**a-d**), data for MCF7 and A549 **(e-f)**. Inhibitor response biomarkers pRB1^S780^ (for CDK4/6 activity) and pMET^Y1234/5^ (for MET activity, determined 48 h following inhibitor addition for b-d **(a)**. Exemplary raw clonogenic data (**b**, **e**), and quantification of clonogenic survival (mean ±SD, n=3 independent repeats) (**c**, **d** and **f**). Palbociclib and crizotinib were used at 450 nM, abemaciclib and foretinib at 225 nM. †††:SI ≥ 0.25 (highly synergistic), ††:0.25 > SI ≥ 0.1 (synergistic), \*\*\*\**p* ≤ 0.0001,1-way ANOVA.

Combinatorial treatment significantly enhanced the reduction in colony outgrowth compared to individual inhibitors (*p* < 0.01, 2-way ANOVA). Identical outcomes were obtained for HCT116 with genomic deletion of the *TP53* gene (*TP53*^−/−^) [33] or isogenic HCT116 with functional *TP53* (*TP53*^WT^), and highly synergistic (SI ≥ 0.25) or synergistic (SI > 0.1) interactions were observed regardless of inhibitor chemotype or combination partner (Fig. 2b-d). Biomarker analysis at 24 h confirmed that inhibitors appropriately modulated their respective targets (Fig. 3a). Thus, pRB1^S780^ had decreased where CDK4/6is were used, while MET auto-phosphorylated on Tyr1234/1235 (pMET^Y1234/5^) had decreased in cells exposed to the METis. As noted previously MET-inhibition did not affect pRB1^S780^ phosphorylation, nor did CDK4/6-inhibition affect the MET-activation state. Cooperativity between MET-inhibition and CDK4/6-inhibition in reducing colony formation capability was also observed in cell lines derived from other cancer types, namely in the oestrogen/progesterone receptor-positive MCF-7 human breast carcinoma-derived cells (MCF-7) and in the KRAS-mutated A549 human lung adenocarcinoma-derived cells (A549) (Fig. 3e and f).

Combined inhibition of MET and CDK4/6 also synergised towards loss of Ki-67 expression (Fig. 4a-d and Supplementary Fig. 2). Loss of Ki-67 is indicative of cell cycle exit [34] and predicts response to CDK4/6-inhibition in preclinical models [7]. Combined inhibition of CDK4/6 and MET for 5 days significantly and cooperatively increased the percentage of HCT116 negative for Ki-67, compared to treatment with single agents—irrespective of *TP53* status (Fig. 4a)—yielding CI values well below 1 across the concentration range tested (Fig. 4b).

**Figure 4.**
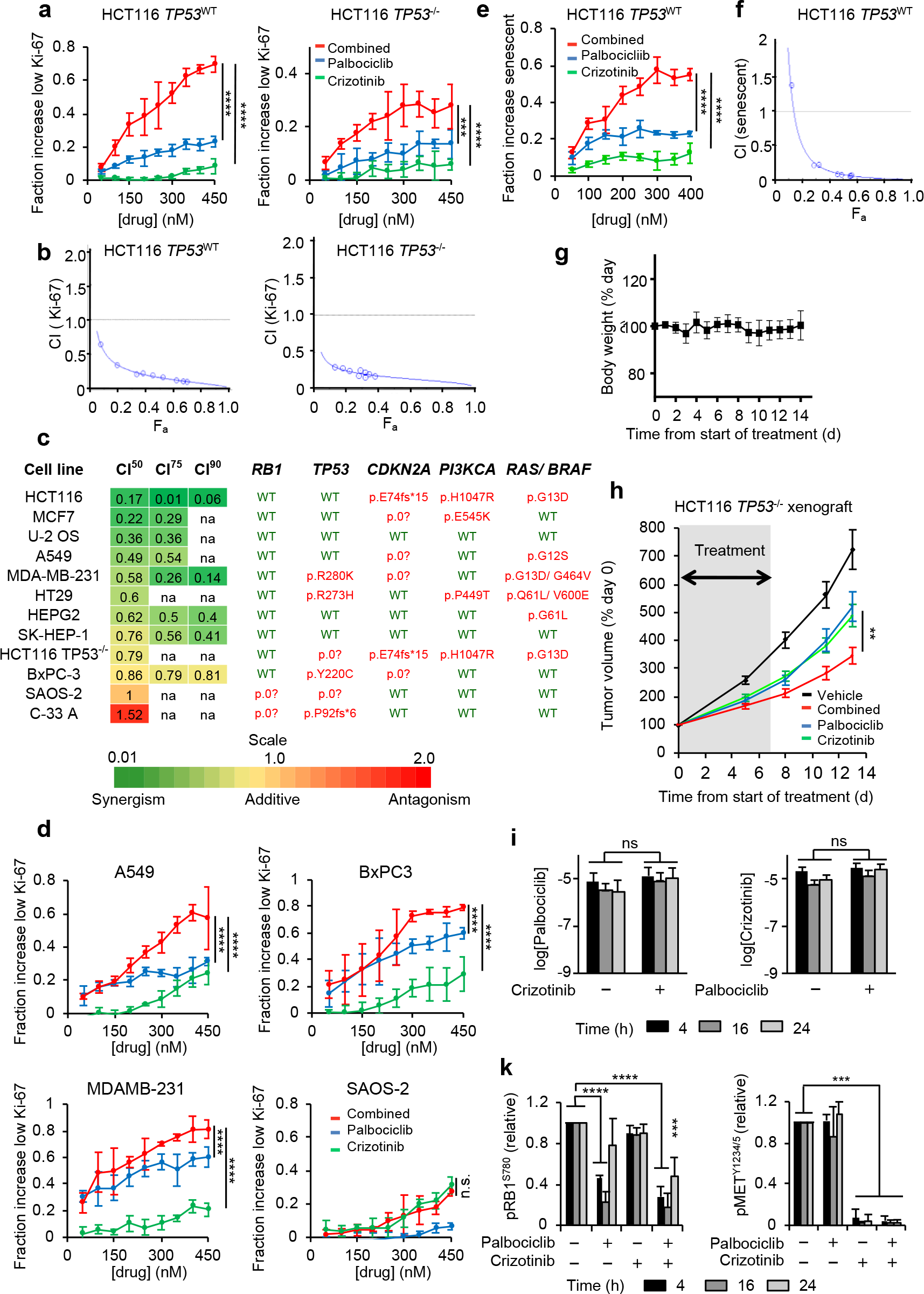
MET and CDK4/6is cooperate to promote cell cycle exit and tumour response *in vivo*. **a** Concentration-effect analysis depicting the fraction of HCT116 with low Ki-67, determined using automated microscopy analysis of cells stained with Ki-67 antibody MIB1. Cells were treated with inhibitors for 96 h. Data represent means ±SD for n=3 independent repeats. **b** CI value plots relating to **a**. **c-d** Effect of CDK4/6 and MET-inhibition on the Ki-67 labeling index across a cancer cell line panel. Cells were treated with palbociclib and/or crizotinib for 96 h. CI value calculation(**c**) and exemplary results (**d**) based on 2 or more independent repeats run in duplicate each are shown. CI^50^, CI^75^ and CI^90^ denote CI values at concentrations with 50, 75 or 90% of cells responding with Ki-67 loss, respectively, na=fractional responses not achievable within the inhibitor concentration range tested. The mutation status of key oncogenic drivers is indicated for each cancer cell line. Data represent mean ±SD for n=2 independent repeats, run in duplicate each. **e-f** Concentration-effect analysis depicting the fractional increase in cells with high SA–β-gal (**e**) and CI value plots (**f**), assessed using automated microscopy of cells reacted with fluorescencent substrate C12FDG. Data represent mean ±SD for n=3 independent repeats, run in triplicate each. **g** Body weight of mice, treated with 100 mg/ kg p.o. crizotinib and 100 mg/kg palbociclib (mean ±SD, n=5). **h** HCT116 tumour xenograft volumes, relative to day 0, in control-and inhibitor-treated mice (mean ±SD, n=10). **i** Concentration of inhibitors in tumour xenograft tissue (mean ±SD, n> 3). **k** Modulation of biomarkers indicative of CDK4/6 (pRB1^S780^) and MET (pMET^Y1234/5^) activity in tumour xenograft tissue (mean ±SD, n = 3). \*\**p*≤ 0.01, \*\*\**p*≤ 0.001, \*\*\*\**p*≤ 0.0001, 2-sided unpaired Student’s t (H, I, K), or 2-way ANOVA comparing the effects size of single agent against that of the combination (**a**, **d**, **e**). (Related to Supplementary Fig. 2 and Supplementary Fig. 3)

Loss of Ki-67 in individual cells correlated with nuclear localisation of the GFP-PSLD reporter in the same cells (Supplementary Fig. 2a and 2b), supporting that reduction of CDK2 activity and loss of Ki-67 are mechanistically linked, Furthermore, Ki-67 positive cells were synergistically reduced in combinations involving different MET and CDK4/6i chemotypes (Supplementary Fig. 2c), indicating that the observed effect is robust and involves on-target MET and CDK4/6-inhibition.

Importantly, MET and CDK4/6-inhibition synergistically reduced Ki-67 expression across cancer cell lines with diverse tissues-of-origin and genetic driver profiles (Fig. 4c and 4d). Concentration-effect analysis confirmed a synergistic interaction for 10 out of 12 lines tested. Notable exceptions were two RB1-mutated lines, the osteosarcoma line SAOS2 and the cervical cancer cell line C-33A, where combination treatment had additive or less than additive effects (Fig. 4c and 4d), consistent with the notion that RB1 is a critical downstream effector of CDK4/6 and that RB1 loss renders cells unresponsive to CDK4/6is. Thus combined MET and CDK4/6-inhibition synergises broadly across cancer histio- and genotypes. The absence of synergism in RB1-mutated backgrounds identifies RB1 activity as a critical component required for this synergistic interaction.

We also assessed if combined inhibition of MET and CDK4/6 enhances or enables senescence by quantifying senescence-associated β-galactosidase (SA-β-gal) in cells. CDK4/6is induce cellular senescence, which is thought to underlie the loss of clonogenic activity they induce [3, 31, 35]. Using C12FDG, a beta-galactosidase substrate with green-fluorescent reaction product suitable for automated quantitative analysis, we observed an overt increase in cells with C_12_FDG fluorescence with characteristic distribution in the perinuclear region following treatment with the inhibitor combination (Supplementary Fig. 3a). Combination-treated cells displayed additional signature changes associated with cellular senescence, including flattened shape, enlarged nuclei and increased cell size (Supplementary Fig. 3a and 3b). Automated quantitative analysis confirmed a significant concentration-dependent increase in the fraction of cells with above baseline perinuclear green fluorescence following combined, compared to single agent treatment with METis and CDK4/6is (Fig. 4e and 3c), with concentration-effect analysis confirming a robust synergistic interaction for the expression of this senescence marker (Supplementary Fig. 3d). Finally, we evaluated cell proliferation activity using cells modified to express the nuclear marker GFP-H2B to track division by time-lapse microscopy (Supplementary Fig. 3e-i). These experiments revealed a synergistic reduction in duplication activity subject to combined inhibition of MET and CDK4/6— irrespective of *TP53* status.

Together these results support the notion that MET-inhibition synergistically increased known cellular responses associated with CDK4/6-inhibition. Conversely, the results support a role for MET in preventing these responses in cells treated with CDK4/6is including responses that predict anti-tumour response to CDK4/6-inhibition *in vivo*.

To test if CDK4/6-inhibition combined with MET-inhibition is a feasible strategy for cancer treatment *in vivo,* we assessed the effect of combined use of crizotinib and palbociclib on the growth of human tumour xenografts in athymic mice. Combined use of 100 mg/ kg (p.o.) crizotinib and palbociclib was well tolerated in the mice (Fig. 4g) yielding effective accumulation of agents within tumour tissue (Fig. 4i) and modulation of pRB1^S780^ and pMET^Y1234/5^ pharmacodynamic biomarkers over a 24 h period (Fig. 4k).

Using this dose we treated mice bearing HCT116 *TP53*^−/−^ tumour xenografts (Fig. 4h) once daily with individual inhibitors or their combination for eight days, then followed them for a further five days, after which time tumours in the control group reached pre-determined size limits. This analysis confirmed superior efficacy of the combination, with statistically significant reduction in tumour burden compared with single agent treatment determined at the end of the observation period (*p* = 0.005 against PD0332991 and *p* = 0.006 against crizotinib, Student’s t tests).

Together, these results provide evidence that MET activity constitutes a resistance mechanism *in vitro* and *in vivo*, broadly reducing the tumour cell response to CDK4/6-inhibition. Importantly, our findings indicate that combined pharmacological inhibition of MET and CDK4/6 is a feasible strategy able to improve tumour growth inhibition in mice, indicating the potential of this combination for clinical use.

### Synergistic inhibition of CDK2 by MET and CDK4/6is involves p21^CIP1^

To identify the mechanism by which MET and CDK4/6-inhibition synergise we characterised activity and composition of the CDK2 complex in inhibitor-treated cells using immunoprecipitation. These experiments confirmed a significant reduction in CDK2 activity subject to combined MET and CDK4/6-inhibition, determined by the ability of the anti-CDK2 immunoprecipitates to yield phosphorylation of substrate (GST-RB 763-928) in vitro (Fig. 5a, 5b, Supplementary Fig. 4b and 4c).

The decrease in CDK2 activity was not accompanied by a decrease in the amount of Cyclin E1 (CCNE1) or a change in the phosphorylation of CDK2 at Tyr15 (pCDK2^Y15^) or Thr160 (pCDK2^T160^), known to confer negative and positive regulation of CDK2 activity, respectively [36] (Fig. 5a). However, there was a significant increase in the amount of the CDK inhibitor protein p21^CIP1^ in precipitates from cells treated with combined MET and CDK4/6i, as compared to single agent-or vehicle-treated cells. Decreased CDK2 activity and increased presence of p21^CIP1^ in anti-CDK2 precipitates were consistently observed in multiple tumour cell lines subject to combined MET and CDK4 inhibition (Fig. 5A, 5C and Supplementary Fig. 4b and 4d). Analysis of input lysates revealed loss of pRB1^S780^ in samples with CDK4/6-inhibition and loss of pMET^Y123/5^ in samples with MET-inhibition (Fig. 5d and Supplementary Fig. 4a), verifying single agent activity and confirming that these proximal biomarkers and pathways are independently modulated by the respective inhibitors regardless of cell background. Essentially identical results were obtained in immunoprecipitations performed using an antibody for the CDK2 activating Cyclin E (Supplementary Fig. 4e-m). Together, these results confirm that reduction of CDK2 activity is a cooperative event caused by combined inhibition of CDK4/6 and MET. They further identify binding of p21^CIP1^ to the CDK2 complex as a potential cause underlying the cooperative interaction of these inhibitors.

**Figure 5.**
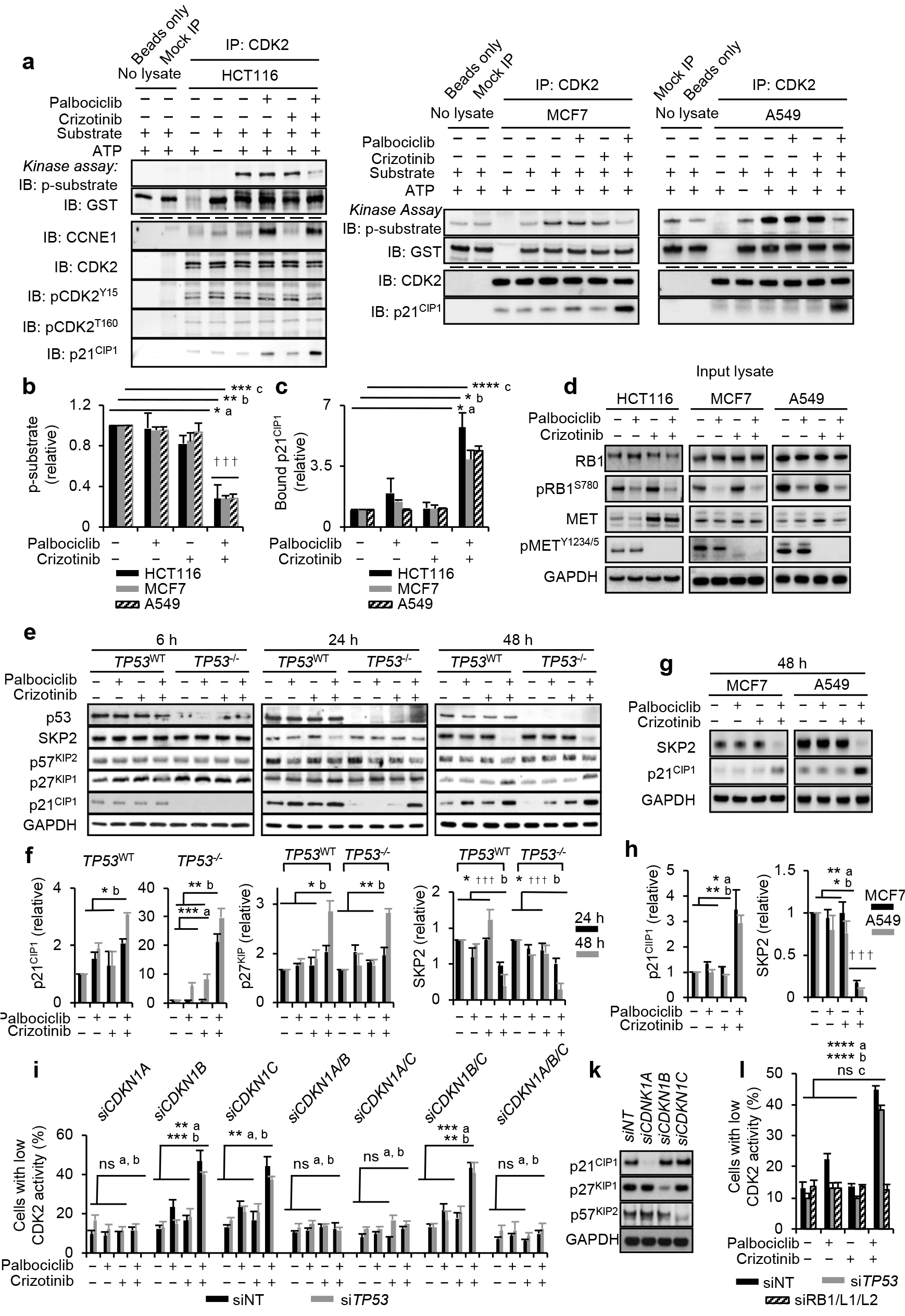
Co-operative control of CDK2 by MET and CDK4/6i involves p21^CIP1^. **a-c** Characterisation of CDK2 complex using anti-CDK2 immunoprecipitation. Data for HCT116, MCF7 and A549 are shown. Cells were treated with 500 nM of inhibitors for 24 h (HCT116) or 48 h (MCF7 and A549). **a** Immunoprecipitation kinase assay, depicting *in vitro* phosphorylated GST-pRB 763-928 substrate (p-substrate) reporting CDK2 activity, and total substrate (GST) (upper), and abundance of co-precipitated p21^CIP1^, total and phosphorylated CDK2 (pCDK2Y15, pCDK2T160), and Cyclin E (CCNE1) in the respective immunoprecipitations (lower). **b** Quantification (mean ±SD, n> 2 independent repeats) of p-substrate and **c** co-precipitated p21^CIP1^ relative to levels in vehicle-treated cells. *p≤ 0.05, \*\**p*≤ 0.01, \*\*\**p*≤ 0.01, \*\*\*\**p*≤ 0.0001, 2-way ANOVA assessing the effect size of single agent with effect size of their combination, ^a^ HCT116, ^b^ MCF7, ^c^ A549. ††† SI ≥ 0.25 (highly synergistic) calculated using mean values. **d** Inhibitor response biomarkers pRB1^S780^ (for CDK4/6 activity) and pMET^Y1234/5^ (for MET activity) for samples analysed in **a-c**. **e-h** Immunoblot analysis assessing levels of CIP/KIP family CDK inhibitors and SKP2 in cell lysates after treatment *of* cell with 500 nM inhibitors. (**e**, **f**) Data for *TP53*^WT^ or *TP53*^−/−^ HCT116, (**g**, **h**) data for MCF7 and A549. (**e**, **g**) exemplary raw data, (**f**, **h**) signal quantification depicting (mean ±SD, n>2 independent repeats). \**p*≤ 0.05, \*\**p*≤ 0.01, \*\*\**p*≤ 0.001, 2-way ANOVA assessing the effect size of single agent crizotinib and palbociclib against effect size of their combination. ^a^24 h, ^b^48 h (**e**, **f**) and aMCF7, bA549 (**g**, **h**). ††† SI ≥ 0.25 (highly synergistic), calculated using mean values. **i** CDK2 activity, assessed using GST-PSLD localization, in HCT116 after transfection with siRNA targeting CIP/KIP family proteins with or without siRNA targeting *TP53*, followed by treatment with 500 nM inhibitors for 24 h (mean ±SD, n=3 independent repeats). ^ns^*p*>0.05, **p≤ 0.01, ***p≤ 0.001, 2-way ANOVA assessing the effect size of single agent against effect size of their combination. aunperturbed (siNT), bTP53-perturbed (siTP53). **k** Immunoblot documenting loss of CIP/KIP family proteins following transfection with siRNA. **l** CDK2 activity, assessed using GST-PSLD localization, in HCT116 after transfection with siRNA targeting RB family proteins or TP53. Data depict mean ±SD for 3 independent repeats, ^ns^*p*> 0.05, \*\*\**p*≤ 0.001, 2-way ANOVA assessing the effect size of single agent treatment against effect size of their combination. ^a^unperturbed (siNT), ^b^TP53-perturbed (siTP53), ^c^RB-perturbed (siRB1/L1/L2). GAPDH (**d**, **e**, **g** and **k**) served as a loading control. (Related to Supplementary Fig. 4)

To assess if the increased association of p21^CIP1^ with the CDK2 complex links to increased p21^CIP1^ abundance, we analysed lysate from treated cells using immunoblotting (Fig. 5e). A progressive increase in p21^CIP1^ abundance was seen between 24 h and 48 h subject to combination treatment (Fig. 5e and 5f). This increase in p21^CIP1^ was apparent in *TP53*^−/−^ HCT116, indicative that the upregulation of p21^CIP1^, a known target transcriptionally activated by TP53, is TP53-independent. We also examined the abundance of the two other members of the CIP/KIP CDK inhibitor family, revealing an increase in p27KIP1 at 48 h, but not at earlier times. No change in the levels of p57KIP2 was observed (Fig. 5e and 5f).

We further assessed the level of the SKP1-cullin-F-box ubiquitin ligase substrate recognition subunit (SKP2), involved in regulating the stability of CIP/KIP family proteins. A clear reduction in SKP2, detectable at 24 h and pronounced at 48 h, was seen following combined inhibition of MET and CDK4/6 compared with vehicle-treated HCT116 (Fig. 5e and 5f). The abundance of SKP2 was not affected following CDK4/6 or MET-inhibition alone. Combined MET and CDK4/6-inhibition leads to upregulation of p21^CIP1^ and loss of SKP2 in multiple cell backgrounds (Fig. 5g and 5h), indicative that this response is broadly observable. Together, these experiments indicate that signalling through MET and CDK4/6 act redundantly to down-regulate the CDK inhibitors p21^CIP1^ and p27^KIP1^, and to up-regulate their regulator SKP2, in turn providing a potential explanation why combined inhibition of MET and CDK4/6 is required for the inhibition of CDK2 and cellular proliferation capacity.

To evaluate if CIP/KIP family members are involved in the synergistic control of CDK2, we depleted the CDK inhibitors alone or in combination using siRNA and, using the localisation of the GFP-PSLD reporter, assessed the effect of this on CDK2 control (Fig. 5i). These experiments positively identified p21^CIP1^ as a critical mechanistic component in the control of CDK2 by combined MET-and CDK4/6-inhibition. Thus, transfection of cells with siRNA targeting *CDKN1A* (which encodes p21^CIP1^) consistently prevented CDK2 control by the combination. siRNA targeting *CDKN1B* (encoding p27^KIP1^) and *CDKN1C* (encoding p57^KIP2^) alone or in combination had no effect, despite evidence that the siRNAs effectively depleted the proteins concerned (Fig. 5k). Similar results were obtained in cells simultaneously transfected with *TP53*-targeted siRNA (Fig. 5i), extending evidence for a critical role of p21^CIP1^, regardless of *TP53* status.

CDK2 regulation by the MET-and CDK4/6i combination was also prevented following loss of RB family proteins (Fig. 5l), consistent with the known resistance to CDK4/6is of cells with functional RB loss and indicative of a critical mechanistic role of RB protein function in the control of CDK2 by combined MET and CDK4/6-inhibition.

### Constitutive FAK activity abolishes cooperation between METis and CDK4/6is

Our siRNA screen identified the FAK family kinases PTK2 and PTK2B, which are known MET effectors, as candidates that facilitate CDK4/6 independent CDK2-activation. Hence we sought evidence if signalling through these kinases is involved in the CDK4/6-independent CDK2-activation by MET and whether additional MET-engaged signalling might play a role. MET and its close relative MST1R connect to multiple effector pathways, amongst them RAS/RAF/ERK, PIK3K/AKT, SRC and STAT3, which they activate in addition to signal transduction involving the FAK/PTK family (Supplementary Fig. 5a).

While effectively blocking MET auto-phosphorylation within the catalytic region (pMET^Y1234/5^) and at the MET multifunctional docking site (pMET^Y1349^) and, further, auto-phosphorylation of MST1R at Tyr1353 (pMST1R^Y1353^) (Supplementary Fig. 5b), treatment of HCT116 with METi alone or combined with CDK4/6i did not affect the activation state of RAS/ERK, PI3K/AKT, SRC and STAT3, indicated by unchanged levels of activated AKT (pAKT^S473^), ERK1/2 (pERK^T202/Y204^), SRC (pSRC^Y416^) and STAT3 (pSTAT3^Y705^), most likely due to activation of these pathways by signalling independent of MET. In contrast, a clear reduction was seen in modifications signifying activation of PTK2 (pPTK2^Y576/577^ and pPTK2^Y925^) and PTK2B (pPTK2B^Y402^), documenting reduced signal transmission through these kinases following MET-inhibition. Together the results identify FAK family activity as critically dependent on MET in HCT116 and loss of FAK family-activation as a key change resulting from MET/MST1R inhibition.

To obtain evidence if FAK family-activation through MET is involved in CDK4/6i-resistant CDK2-activation, we generated HCT116 expressing a membrane-targeted, constitutively active PTK2 variant, CD2-PTK2 (Fig. 6a). Expression of CD2-PTK2 did not affect the ability of MET-inhibition to block MET receptor activity, or the ability of CDK4/6-inhibition to block RB1 phosphorylation (Fig. 6a). However, expression of CD2-PTK2 permitted sustained, METi-resistant PTK2-activation, indicated by the presence of pY576/577 phosphorylated CD2-PTK2 (Fig. 6a). Significantly, in cells expressing CD2-PTK2 the activity of CDK2 in anti-CDK2 immunoprecipitates was unaffected by combined inhibition of CDK4/6 and MET (Fig. 6b and 6c). Furthermore, the increased association of p21^CIP1^ following combined CDK4/6 and MET-inhibition was abolished (Fig. 6b and 6d and Supplementary Fig. 5c - 5f).

**Figure 6.**
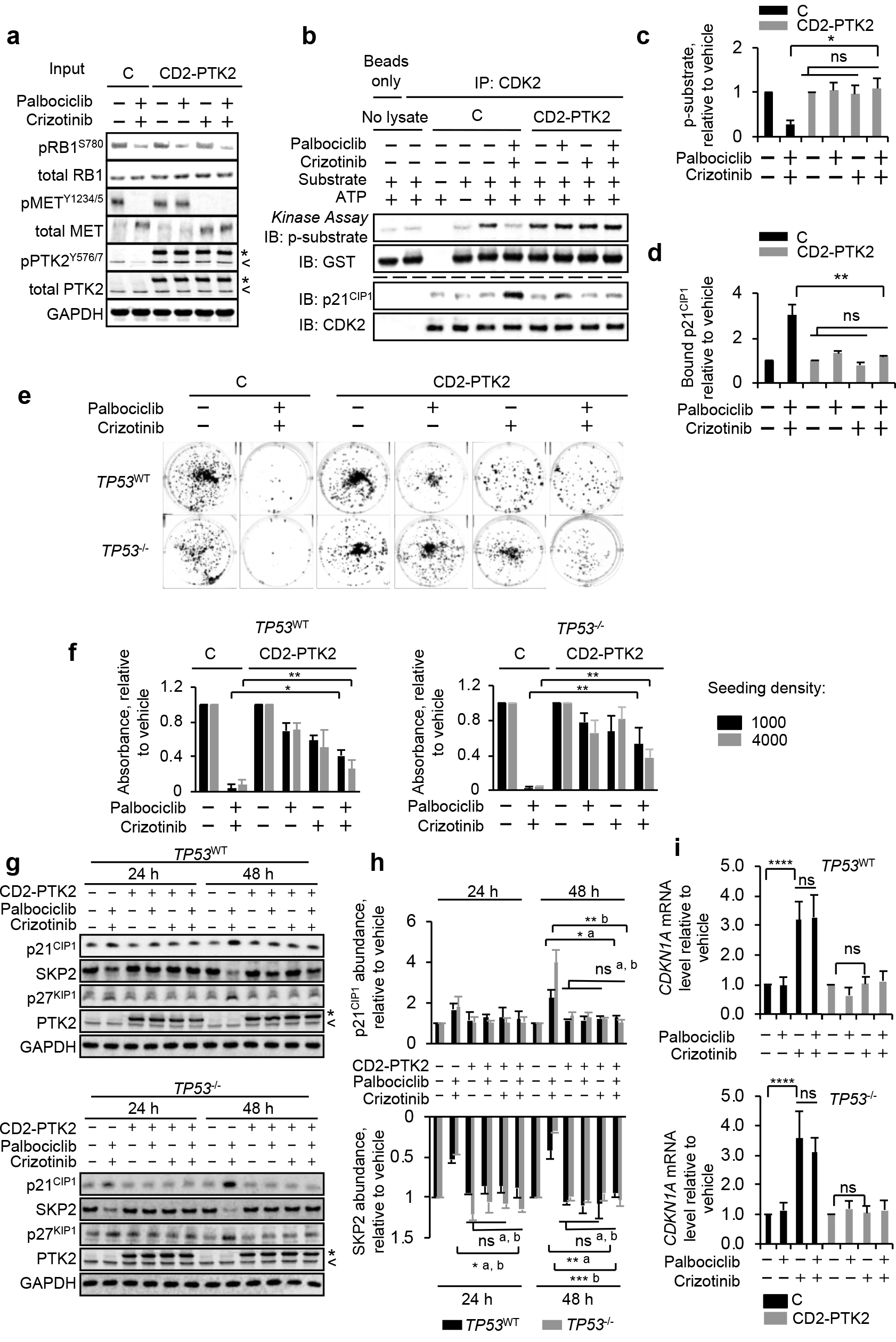
Constitutive PTK2 activity impairs co-operation between MET and CDK4/6is. **a-d** Characterisation of CDK2 complexes in HCT116 expressing constitutively active CD2-PTK2. **a** Immunoblot of input lysate documenting CD2-PTK2 expression and modulation of drug response biomarkers pRB1^S780^ (for CDK4/6 activity) and pMET^Y1234/5^ (for MET activity). C denotes control cells without CD2-PTK2 expression, < identifies signal for cell-intrinsic PTK2, * identifies CD2-PTK2. **b** Immunoprecipitation kinase assay. Abundance of *in vitro* phosphorylated GST-pRB 763-928 substrate (p-substrate) reflecting CDK2 activity and total substrate (upper), abundance of p21^CIP1^ and CDK2 (lower) in the respective immunoprecipitations. (**c**) Mean abundance of p-substrate and (**d**) mean abundance of co-precipitated p21^CIP1^ relative to vehicle-treated cells. Data (**c**, **d**) represent mean ±range for 2 independent repeats. Cells were treated for 24 h using 500 nM of each inhibitor. \**p*≤ 0.05, \*\**p*≤ 0.01, 2-way ANOVA assessing the effect size of combination treatment between control and CD2-PTK2 expressing cells and ^ns^*p>*0.05, 2-way ANOVA assessing the effect size of single agent against effect size of their combination in cells expressing CD-PTK2. **e-f** Clonogenic survival of cells expressing constitutively active CD2-PTK2. (**e**) Exemplary raw images and (**f**) quantitative assessment representing mean values ±SD for 3 independent repeats. Cells were treated with 500 nM inhibitors for 120 h. \**p*≤ 0.05, \*\**p*≤ 0.01, 2-way ANOVA assessing effect size of combination treatment between controls and CD2-PTK2 expressing cells. **g-h** Abundance of CIP/KIP family proteins and SKP2 in HCT116 with constitutively active CD2-PTK2. Cells were treated with 500 nM inhibitors. (**g**) Exemplary raw data and (**h**) quantitative assessment representing mean ± range of 2 independent repeats. \**p*≤ 0.05, \*\**p*≤ 0.01, 2-way ANOVA assessing effect size of combination treatment between controls and CD2-PTK2 expressing cells and ^ns^*p* > 0.05, 2-way ANOVA assessing effect size of single agent against effect size of their combination in cells expressing CD-PTK2. **i** RT/qPCR assessing *CDKN1A* mRNA levels. Cells were treated with 500 nM inhibitors for 24 h. Data are mean values ±SD of 3 independent repeats; \*\*\*\**p*≤ 0.0001, ns*p*> 0.05, 1-way ANOVA (Related to Supplementary Fig. 5)

Notably, expression of CD2-FAK reduced the ability of combined CDK4/6 and MET-inhibition to decrease colony formation in HCT116 (Fig. 6e and 6e), irrespective of TP53 status. It also reduced the increase in p21^CIP1^ and the loss of SKP2, associated with combined MET and CDK4/6-inhibition (Fig. 6g and 6h). Together these results indicate that reduction of FAK activity is key to limiting CDK2 and clonogenic activity following from combined MET and CDK4/6-inhibition in cells.

The finding that combined inhibition of the MET/FAK and CDK4/6 axis can promote p21^CIP1^ accumulation in HCT116 *TP53*^−/−^ implies a mechanism for *TP53*-independent generation of p21^CIP1^, conferred by inhibition of the MET/FAK axis. Consistent with this, and regardless of TP53 status, we observed a significant increase in CDKN1A mRNA (encoding p21^CIP1^) following combined CDK4/6 and MET-inhibition compared to control or CDK4/6i treated cells, (Fig. 6i), which was abolished by the expression of constitutively active PTK2. Notably, the level of *CDKN1A* mRNA increased in cells treated with METi alone, indicating that *CDKN1A* transcription is suppressed by signalling through MET/FAK, independent of CDK4/6. Together, these experiments identify FAK family activity as a critical effector downstream of MET involved in preserving CDK2 activity and clonogenic potential in cells exposed to CDK4/6i. They further suggest TP53-independent *CDKN1A* transcript accumulation as a candidate mechanism by which inhibition of MET/FAK promotes CDK4/6i-resistant CDK2-activation and cell cycle activity.

### SKP2 is the critical common target engaged by MET and CDK4/6

In addition to the increase in p21^CIP1^ our earlier results showed a loss of SKP2 following combined CDK4/6 and MET-inhibition (Fig. 5e-h). SKP2 mediates the degradation of p21^CIP1^ [37] and itself is targeted for degradation involving RB1 [15, 16]. Therefore, SKP2 loss may be the consequence of sustained RB1 activity, following from effective control of CDK2 activity through p21^CIP1^. And SKP2 loss may stabilise p21^CIP1^, synthesised from transcript that accumulates as a consequence of inhibition of the MET/FAK axis.

To test if this model explains how METis and CDK4/6is synergise, we used siRNA or inhibitors to ablate relevant components and assessed the effect of this on SKP2 loss and p21^CIP1^ accumulation (Fig. 7). Consistent with the model ablation of RB proteins abolished SKP2 loss and p21^CIP1^ accumulation following combined MET and CDK4/6-inhibition (Fig. 7a and 7b). Also consistent with this model, PTK2 ablation, or pharmacological inhibition of FAK family activity using defactinib, were sufficient to trigger p21^CIP1^ accumulation but insufficient to cause SKP2 loss, for which additional inhibition of CDK4/6 was required (Fig. 7c and 7d).

**Figure 7.**
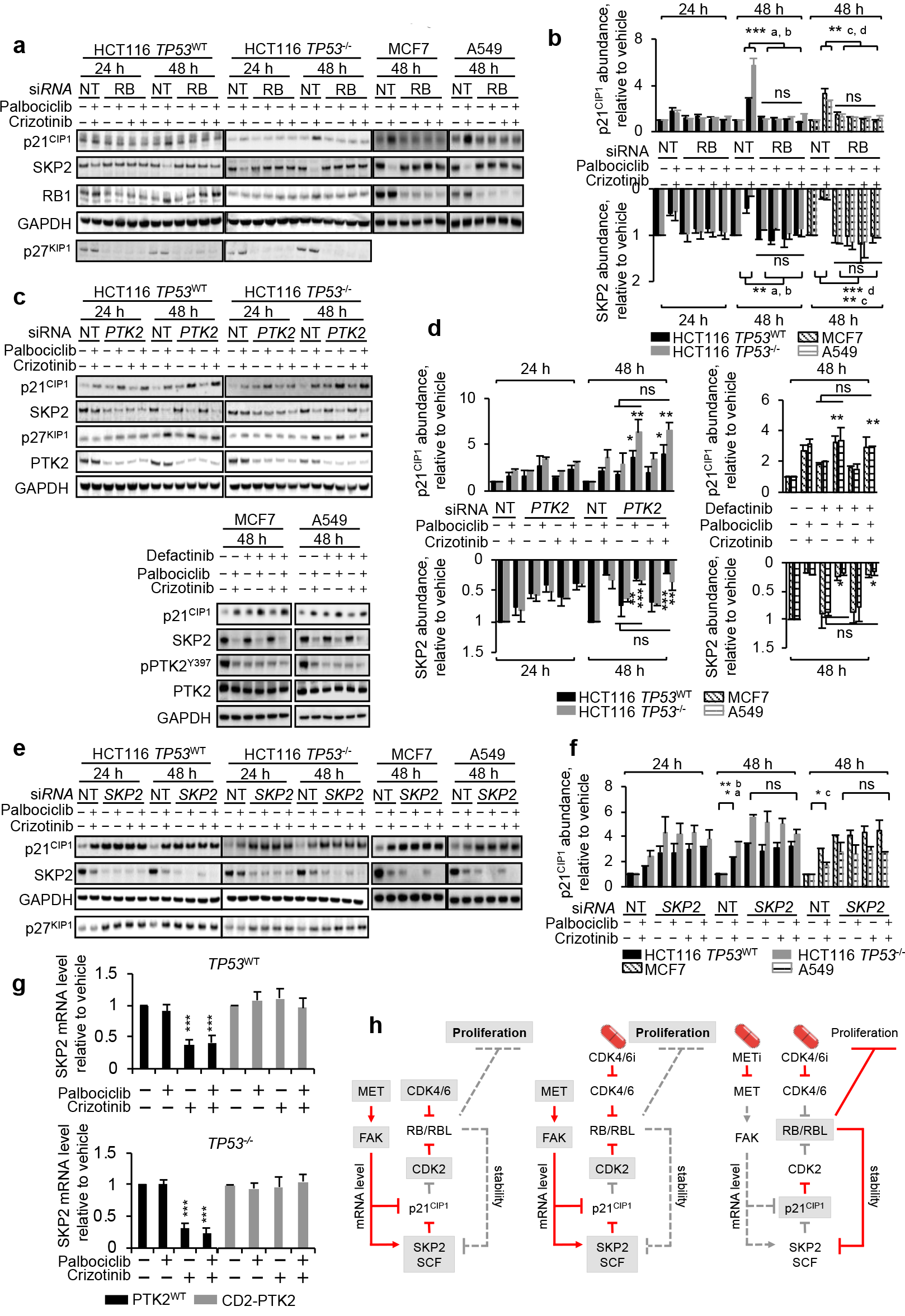
SKP2 is the critical function engaged by MET and CDK4/6. **a-f** Levels of p21^CIP1^, p27^KIP1^, SKP2 protein in cells treated with siRNA against *RB* (**a**, **b**), siRNA against *PTK2* or the FAK inhibitor defactinib (900nM) (**c**, **d**), or siRNA targeting *SKP2* (**e**, **f**). Data for *TP53*^WT^ and *TP53*^−/−^ HCT116, MCF7 and A549 are shown. Cells were treated with 500 nM METi crizotinib and/ or CDK4/6i palbociclib. Representative raw data (**a**, **c**, **e**), and mean ± range for 2 independent repeats (**b**, **d**, **f**). 2-way ANOVA comparing in (**b**) the effect size of the combination in control and RB-depleted cells (\*\**p*≤ 0.01, \*\*\**p*≤ 0.001) or the effect size of single agent with that of their combination in RB-depleted cells (^ns^*p* > 0.05), in (**d**) the effect size of single agent palbociclib with that of the combination (^ns^*p* > 0.05). 1-way ANOVA assessing in (**d**) the effect of treatment compared to vehicle (\**p*≤ 0.05, \*\**p*≤ 0.01, ***p≤ 0.001), in (**f**) the effect of vehicle vs. that of the combination. ^a^ *TP53*^WT^ +/+, ^b^ *TP53*^−/−^, ^c^ MCF7, ^d^ A549, **g** RT/qPCR assessing *CDKN1A* mRNA in HCT116 treated with 500 nM inhibitors for 24 h. Data are mean values ±SD for 3 independent repeats; \*\**p*≤ 0.01, 1-way ANOVA comparing the effect of vehicle vs. treatment. **h** Model, detailing consequence of combined CDK4/6-and MET-inhibition. Active nodes underplayed in grey; active signaling in red.(Related to Supplementary Fig. 6)

Unexpectedly, SKP2 loss (Fig. 7e and 7f), predicted to require MET-inhibition in order to yield upregulation of p21^CIP1^, yielded p21^CIP1^ accumulation that was not further enhanced by MET-inhibition. This result indicates that p21^CIP1^ transcript up-regulation caused by MET-inhibition may not be essential for p21^CIP1^ accumulation, at least under conditions where SKP2 is absent at the outset, and could indicate that MET/FAK inhibition could have involvement in promoting SKP2 loss, independent from the degradation that it may enable by cooperating with CDK4/6-inhibition towards RB1-activation. Consistent with this prediction we found that MET-inhibition, in addition to increasing transcript levels for p21^CIP1^, suppressed transcript levels for SKP2, which like the increase in p21^CIP1^ transcript was counteracted by constitutively active PTK2 (Fig. 7g). Also consistent with this prediction we found that SKP2 loss enabled palbociclib to control CDK2-activation without additional MET-inhibition (Supplementary Fig. 6).

Together these data support a mechanistic model (Fig. 7h) in which SKP2 acts as a common target engaged by MET and CDK4/6 and proposes FAK-driven SKP2—and potentially p21^CIP1^ transcript regulation—as mechanistic events through which the MET/FAK axis confers refractory response to CDK4/6-inhibition, and through which MET-inhibition synergises with CDK4/6-inhibition to deliver increased anticancer activity.

## Discussion

Widespread recognition exists that CDK2-activation is associated with acquired resistance of cancer cells to CDK4/6-targeting cancer therapeutics [21–23] and also that CDK2 supports CDK4/6-independent proliferation during organismal development [20]. However, the molecular determinants that permit activation of CDK2 in these contexts have not been systematically sought. CDK4/6-selective inhibitors are now showing considerable promise in patients with estrogen receptor-positive breast cancer, yet there is clear need to identify functions that drive resistance in treatments involving these agents and that preclude their broader use in other cancer types. Here we report that activation of the MET/FAK signalling axis leads to CDK4/6-independent CDK2 activation, and constitutes a broadly applicable druggable means to improve the response of cancers to CDK4/6-targeted therapies.

MET is widely expressed in epithelial and endothelial cells, including cancers derived from these tissues, and may be activated in cancer cells through mutation. However, more frequently MET signalling is activated by the hepatocyte growth factor/ scatter factor, HGF, produced by adjacent mesenchymal tissues, including stromal components of cancers [38–40].

We identified the MET/FAK axis based on a functional genetic screen using CDK2 activity as a mechanism-based endpoint. Apart from the MET/FAK axis, which we mechanistically explore in our work, the screen yielded other hits without known links to MET and/or FAK signalling, including ontology noted previously for synergistic interaction with catalytic inhibition of CDK4/6 such as RPS6KA6 and BRAF, inhibitors of which increase the response of cancer cells to CDK4/6-inhibition [2, 21, 41]. Unexpectedly, our screen indicates that *TP53* status affects the spectrum of molecular functions required to support CDK4/6 independent CDK2-activation, with requirement of certain functions, including e.g. RPS6KA6, confined to *TP53* normal backgrounds. While further validation is required, these observations suggest that the *TP53* status could determine efficacy of certain combinations.

In addition to MET, the screen identified FAK family kinases, that our subsequent work validates as a critical component by which MET elicits CDK4/6-independent CDK2-activation. Our experiments involving a constitutively-active form of the FAK family kinase PTK2 demonstrate that FAK activity is sufficient to elicit significant CDK4/6i tolerance. In the experimental models examined, MET scores as a key determinant responsible for FAK family-activation, as indicated by the robust reduction of activated FAK species pPTK2^Y576/577^, pPTK2^Y925^ and pPTK2B^Y402^ following treatment with METi. However, FAK family kinases can be activated by other routes including MET-unrelated tyrosine kinase receptors and their activators, and by extracellular matrix signalling involving integrins [42]. The broad range of signalling able to engage FAK highlights the possibility that CDK4/6i resistance involving FAK could be caused in cancer patients by events other than MET-activation.

Our work indicates that sustained expression of the ubiquitin ligase subunit SKP2, which promotes the degradation of including p21^CIP1^ as well as that of other CIP/KIP CDK inhibitors, is a key mechanism by which MET/FAK supports activation of CDK2. While we currently do not know how exactly MET/FAK signalling effects SKP2 expression, our experiments show that signalling through this axis increases the steady state level of SKP2 mRNA. The involvement of FAK in the regulation of SKP2 and p21^CIP1^ was previously noted. For example, inhibition of FAK through enforced expression of FAK-related non-kinase (FRNK) or dominant negative FAK (Tyr397Phe) reduced the expression of SKP2 protein in human and rat endothelial cells [43, 44]. Furthermore, inhibition of FAK by FRNK, FAK (Tyr397Phe) or small molecule inhibitors resulted in elevated p21^CIP1^ transcript and protein in normal human fibroblasts, smooth muscle cells or glioblastoma-derived cells [43, 45, 46]. These results independently support our observations that FAK family kinases regulate SKP2 and the CIP/KIP inhibitor p21^CIP1^.

Our work indicates that MET-inhibition significantly enhances therapeutic inhibition of growth by CDK4/6is in preclinical models of human cancer. Furthermore, our results identify FAKs as key downstream components in this context. The recognised ability of cancer-relevant, MET-independent signalling routes to activate FAKs make this kinase family, or druggable targets downstream of it, potentially more attractive therapeutic targets than MET for use in combination with CDK4/6-inhibition. FAK family kinases are a recognised drug target in cancer and inhibitors targeted to the ATP-binding pocket of their kinase domain have entered clinical trials, albeit, with limited single-agent efficacy in patients thus far [47– 49]. Our results suggest opportunity for use of FAK inhibitors in conjunction with CDK4/6is as a potentially powerful approach to improve the outcome for patients treated with CDK4/6-targeted therapies or to expand the current, approved indications through mechanism-based targeted combinations.

## Methods

### Cell culture, chemicals and antibodies

HCT116 *TP53*^−/−^ and isogenic HCT116 *TP53*^wt^ cells were provided by the Vogelstein laboratory (John Hopkins University, Baltimore, MD). Other cell lines were acquired from the American Type Culture Collection (ATCC). HCT116-PSLD are described in [26]. Cells expressing CD2-PTK2 were constructed by lentiviral transduction using pLV-neo-CD2-FAK [50]. Inhibitors used were purchased from Selleck Chemicals. Antibodies and siRNAs used are summarized in supplementary materials.

### RNAi screens

Screens used the kinase-covering component of the Dharmacon siGENOME SMARTpool™ library. Library pools were mixed at equimolar ratio with SMARTpool™ oligonucleotides targeting *TP53* or non-targeting oligonucleotide, then reverse transfected at a combined concentration of 20nM into HCT116-PSLD, seeded in 96-well plates. Transfected cells were incubated for 24 h prior to treatment with CDK4/6i palbociclib (450nM) or vehicle for 24 h. Plates were fixed in 4% formaldehyde, then stained with Hoechst 33342 DNA dye and imaged using an INCell Analyzer 3000 (GE Healthcare) or an Opera (Perkin-Elmer) high-content imager platform. Data for a minimum of 2000 cells per condition were collected. Data were processed using CellProfiler open-source image analysis software [51].

### In vivo human tumour xenograft studies

All animal work was carried out under UK Home Office regulations in accordance with the Animals (Scientific Procedures) Act 1986 and according to United Kingdom Co-ordinating Committee on Cancer Research guidelines for animal experimentation [52] with local ethical approval. For therapy studies, mice were treated daily for a continuous period, followed by observation until tumour size in the control group approached pre-determined humane size limits. For pharmacodynamic studies, tumour samples were collected at 24 h post-administration.

Further method details are provided in the supplementary text.

## Acknowledgments

The authors thank Nicky Evans for editorial help with writing this manuscript, and Albert Hallsworth and Melanie Valenti for assisting with the preclinical work. We thank the Vogelstein laboratory for providing isogenic *TP53*^−/−^ and *TP53*^WT^ HCT116 human colorectal cancer cells. Financial support: The work was supported by grants from Cancer Research UK (ref. C309/A11566, ref. C368/A6743 and ref. A368/A7990 (PW), ref. C423/A1421 and ref. C423/A15043 (SM)) and the World Cancer Research Fund (WCRF) (ref.12-1280). CZ was supported by a Wellcome Trust studentship (ref. 094885/Z/10/Z). PW is a Cancer Research UK Life Fellow. The funders had no role in design of the study, the collection, analysis, and interpretation of the data the writing of the manuscript or the decision to submit the manuscript for publication.

## Author contribution

S.M., P.W., P.C. and S.R.S. supervised the project. C.Z. and S.M. wrote the manuscript; C.Z., S.R.S, S.M., F.R. and S.E. designed experiments; B.A-L. supervised and C.Z. performed the bioinformatics analysis; R.K. and J.F. supported the high content data acquisition and data extraction; S.E. designed the xenograft studies, performed by A.D.H.B.; C.Z. performed the majority of experiments; A.H. performed assessment and F.R. analysed outcome of the pharmacokinetics analysis; M.E. performed the Ki67 based response analysis; C.Z, S.R.S., S.E., F.R. and S.M. analyzed data.

