## supplementary text and figures and tables for "Signalling involving MET and FAK supports cell division independent of the activity of the cell cycle-regulating CDK4/6 kinases"

**Supplementary Material and Methods**

**Cell culture, chemicals and antibodies:** HCT116 *TP53*$^{-/-}$ and isogenic HCT116 *TP53*$^{wt}$ cells were provided by the Vogelstein laboratory (John Hopkins University, Baltimore, MD). All other cell lines were acquired from the American Type Culture Collection (ATCC). HCT116 stably expressing the GFP-PSLD reporter were described [1]. Cells expressing CD2-PTK2 were constructed by lentiviral transduction using pLV-neo-CD2-FAK [2]. All inhibitors used were purchased from Selleck Chemicals. Antibodies and siRNAs used are summarized in supplementary materials. Antibodies for AKT (RRID:AB_329827), pAKT$^{S473}$(RRID:AB_331161), ERK (RRID:AB_390780), p27$^{KP1}$ (RRID:AB_2077850), pMET$^{Y1234/5}$ (RRID:AB_10691464), pPTK2$^{TYR397}$ (RRID:AB_10891442), pPTK2$^{Y576/7}$ (RRID:AB_10891442), pPTK2$^{Y925}$ (RRID:AB_2253227), PYK2 (RRID:AB_2174097), pPYK2$^{Y404}$ (RRID:AB_2300530), pSRC$^{Y416}$ (RRID:AB_10860245), RB1 (RRID:AB_10696874) and pSTAT3$^{Y705}$ (RRID:AB_2491009) were purchased from Cell Signalling Technology. Antibodies against CDK2 (RRID:AB_2078401), pCDK2$^{Y15}$ (RRID:AB_1310069), pCDK2$^{T160}$ (RRID:AB_869087), Cyclin E1 (RRID:AB_10973355) used in immunoblotting, GAPDH (RRID:AB_2278693), p53 (RRID:AB_471101), p57$^{KIP2}$ (RRID:AB_1267271), pRB1$^{S780}$ (RRID:AB_777635), pMST1R$^{Y1353}$ (RRID:AB_10973056), SRC (RRID:AB_870739) and STAT3 (RRID:AB_10901752) were purchased from Abcam. Antibodies against Cyclin E1 (RRID:AB_627362) used for immunoprecipitation, RBL1 (RRID:AB_628058), RBL2 (RRID:AB_632093), MST1R (RRID:AB_677390) were purchased from Santa Cruz. Antibodies against p21$^{CIP1}$ (RRID:AB_1793044), pERK1/2$^{Y202/4}$ (RRID:AB_259347), Ki-67 (RRID:AB_2631211), and pRB1$^{S807}$ (RRID:AB_628602) were purchased from Upstate, Sigma-Aldrich, Dako and Abnova respectively. Serum against Gluthatione-S-Transferase (GST) was raised in rabbits using recombinant GST protein, purified from E., coli as immunogen.

**Cell culture:** Cells were grown in polystyrene flasks (BD Biosciences) in Dulbecco's Modified Eagle Medium DMEM medium and 5% $CO_2$ at 37 °C, with the exception of HT29, U-2OS and SAOS-2 cells (McCoy’s 5a medium) and BXPC-3 cells (RPMI-1640 medium). All media were supplemented with 10% FBS. All cell lines used were confirmed *Mycoplasma* free using PCR combined with culture-based testing. Cell lines were obtained authenticated from ATCC. Where cell lines were obtained from another source, or were subjected to genetic modification, authentication was performed using STR-based methodology. siRNA transfections were performed using HiPerFect transfection agent (Qiagen). DNA transfections used FuGENE 6

(Promega). siRNAs for *SKP2* (M-003324-04), *TP53* (M-003329-03), *CDKN1A* (M-003471-00), *CDKN1B* (M-003472-00), *CDKN1C* (M-003244-03), *RB1* (L-003296-02), *RBL1* (M-003298-02), *RBL2* (M-003299-03), *MET* (M-003156-02), *MST1R* (M-003157-04), *PTK2* (M-003164-02), *PTK2B* (M-003165-03), *SRC* (M-003175-03), *ABL1* (M-003100-02) and *STAT3* (M-003544-02) were obtained from Dharmacon. Non-targeting control siRNA (AllStars) and siRNA against *CDK2* (Hs_CDK2_11, Hs_CDK2_12) were obtained from Qiagen.

**RNAi screen:** Screens used the kinase-covering component of the Dharmacon siGENOME SMARTpool™ library. Library pools were mixed at equimolar ratio with SMARTpool™ oligonucleotides targeting *TP53* or non-targeting oligonucleotide, then reverse transfected at a combined concentration of 20nM into HCT116-PSLD, seeded into triplicate wells of opaque, tissue culture treated, 96-well plates with transparent base (Packard View plates, Perkin Elmer). Transfected cells were incubated for 24 h prior to treatment with CDK4/6 inhibitor palbociclib (450nM) or vehicle (DMSO) for 24 h. Plates were fixed in 4% formaldehyde for 10 minutes at room temperature, Prior to imaging the fixed cells were permeablized in TBS buffer (25 mM Tris, 140 mM NaCl, pH 7.5) containing 0.1% Triton X-100 supplemented with 1 μg/ ml Hoechst 33342 DNA dye and imaged using an INCell Analyzer 3000 (GE Healthcare) or an Opera (Perkin-Elmer) high-content imager platform. Data were processed using CellProfiler open-source image analysis software [3] as described in [4]. A custom perl script, described in [4] was used to compute the percentage of imaged cells with nuc/cyto fluorescence ratio of > 1.5 in each well.

**Immunoblot analysis:** Cells were lysed in ELB buffer (50 nM HEPES, 250 nM NaCl, 0.5% Triton X-100, 1 mM EDTA, pH 7.4) containing complete protease inhibitor cocktail (Roche) and phosphatase inhibitors (1 mM NaF, 10 mM β-glycerophosphate and 4 mM $Na_3VO_4$). Protein concentrations were measured by BCA protein assay (Pierce). Samples were separated on SDS polyacrylamide gels and transferred to an Immobilon-FL membrane (Millipore), before incubation with primary antibodies overnight at 4 °C. Membranes were washed with TBST (25 mM Tris, 140 mM NaCl, 0.1% Tween-20, pH 7.5) and probed with IRDye 680- or IRDye 800CW-conjugated secondary antibodies (LI-COR) for 1 h at room temperature. The imaging and quantification of protein bands was carried out using an Odyssey CL (LI-COR) infrared imaging system.

**Immunoprecipitation and *in vitro* kinase assay.** Cells were lysed in ice-cold immunoprecipitation (IP) buffer (50 mM HEPES, 150 mM NaCl, 1 mM EDTA, 2.5 mM EGTA, 50 mM NaF, 20 mM β-glycerophosphate, 4 mM $Na_3VO_4$, 1 mM DTT and 0.1% Tween-20, pH

8), lysates were cleared by centrifugations at 10,000 x g at 4 °C. Cleared supernatants were mixed with 50 µl of Dynabead coupled protein A (Life Technologies) bound with 2 µg of IgG. Beads were incubated with rotation for 20 min at 4 °C, then washed three times in IP buffer. Precipitates were analysed using SDS polyacrylamide gel electrophoresis followed by immunoblotting or were used in vitro kinase activity assays. Kinase activity assays were performed as previously described [5], using as a substrate GST-RB1$^{763-928}$ recombinant protein (500 ng/reaction), purified as described [6], in 20 µl kinase reaction buffer (25 mM HEPES, 0.1% β-mercaptoethanol, 0.1 mM EGTA, 25 mM $MgCl_2$ and 400 µM ATP, pH 7.9) added directly to the beads. The mixture was incubated for 10 min at 30 °C, reaction products were separated on SDS polyacrylamide gels and subjected to immunoblot analysis using antibody directed against the proline-directed phosphorylation site Ser807 of human RB1 (anti-pRB1$^{S807}$) to quantify substrate phosphorylation and rabbit anti-GST antiserum, to visualise substrate present in each reaction.

**Cell-based immunostaining assay:** Antibody staining was performed in 96 well plates as in [1]**.** Cells grown in 96-well plates were fixed with 4% formalin for 10 minutes, permeablized in TBS (25 mM Tris, 140 mM NaCl, pH 7.5) supplemented with 0.1% Triton X-100 and then washed in TBS containing 0.1% Tween 20 (TBST) and 5% skimmed milk. Primary antibodies were diluted to 1:250 in TBST. Cells were incubated with diluted antibodies overnight at 4 ºC, prior to washing in TBST buffer. Cells were subsequently probed with Alexafluor-647®-coupled secondary antibodies containing 1 µg/ mL Hoechst 33342. Cells were imaged as described preciously [4]. A custom script, described in [4], was used to calculate the percentage of cells with a nuclear fluorescence intensity above threshold.

**Detection of senescence-associated β-Galactosidase activity:** Detection of SA-β-Gal activity using with $C_{12}$FDG staining (Life Technologies) was carried out as previously described [7]. Fluorescence intensity data within the perinuclear region of individual cells was quantified using CellProfiler free open-source image analysis software (http://cellprofiler.org/). Images of Hoechst 33342 fluorescence were used to generate a nuclear mask with which each imaged cell was located and subsequently measured for perinuclear staining in corresponding $C_{12}$FDG fluorescence images. Thresholds to gate for cells exhibiting SA-β-Gal positivity above background were determined using histogram plots generated using the 'R' free statistical software (https://www.r-project.org). A custom perl script, described in [4], was used to calculate the percentage of cells with a perinulcear $C_{12}$FDG intensity above threshold.

**Time-lapse microscopy.** Time-lapse observations were carried out in cells expressing H2B-GFP using a IncuCyte™ live cell analysis system (Essen Bioscience). Cells were seeded into

96-well plates 24 h prior to treatment, and followed by fluorescence imaging every two hours at 37 °C with 5% $CO_2$. Data were collected from a minimum of three parallel wells per condition.

**Clonogenic assay.** Cells seeded into 6-well plates were treated with drug(s) for 5 days, then detached using trypsin. Cells were counted and reseeded at fixed cell numbers into duplicate wells of 6-well plates into drug-free medium. After 14 days cells were fixed with 4% formaldehyde, and stained with 0.5% crystal violet dye dissolved in 25% methanol. Excess dye was thoroughly removed by repeat washing with tap water. Digital images were generated using a PC desktop scanner, then bound crystal violet dye was eluted using 30% v/v acetic acid, 30% v/v DMSO, 1% SDS and photometrically quantified at 590 nm absorbance.

***In vivo human* xenograft studies:** Compounds were dissolved in sodium lactate pH 4.0 (palbociclib) or sterile saline (crizotinib) and administered at 100 mg/ kg once daily p.o.. Six to seven week- old female NCr-*Foxn1*$^{nu}$ mice were inoculated s.c. into the right flank with 3 x $10^6$ HCT116 *TP53*$^{-/-}$ cells. Treatment was initiated when tumour grafts reached 5 to 6 mm in diameter. Tumour size was measured twice per week using callipers and volumes were calculated using the formula 1/2 (length (mm)) × (width (mm)). Pharmacokinetic studies used a single dose of 100 mg/ kg palbociclib and/or crizotinib, administered p.o. Plasma and tumour samples were collected at 4, 16 and 24 h post-administration, and analysed as described previously [8]. For pharmacodynamic studies, tumour samples were collected at 24 h post-administration.

**Statistical Analyses.** Statistical assessments used two-tailed unpaired Student's t-test, 1-ANOVA and 2-way ANOVA. Calculations were performed using GraphPad 7.0. Calculation of p-values for pathway enrichment analysis, measuring the likelihood that the intersection between the input genes and a particular network was obtained by chance, were carried out using MetaCore™ as described in the user manual. All data are expressed as normalized means and standard deviation (SD) or standard error (SE) from multiple independent measurements, as stated. Z-scores, describing the distance from the target mean to the population mean in units of standard error, were computed using Z-test statistics in EXCEL. Gene clustering was performed using the heatmap function within 'R' open source software (https://www.r-project.org). Tests for interaction between target knockdown and treatment were performed as previously described [1]. The degree (index) of interaction (SI) was calculated by subtracting the observed combined effect of treatments from the product of the observed individual effects.

### Supplementary Figures and Legends

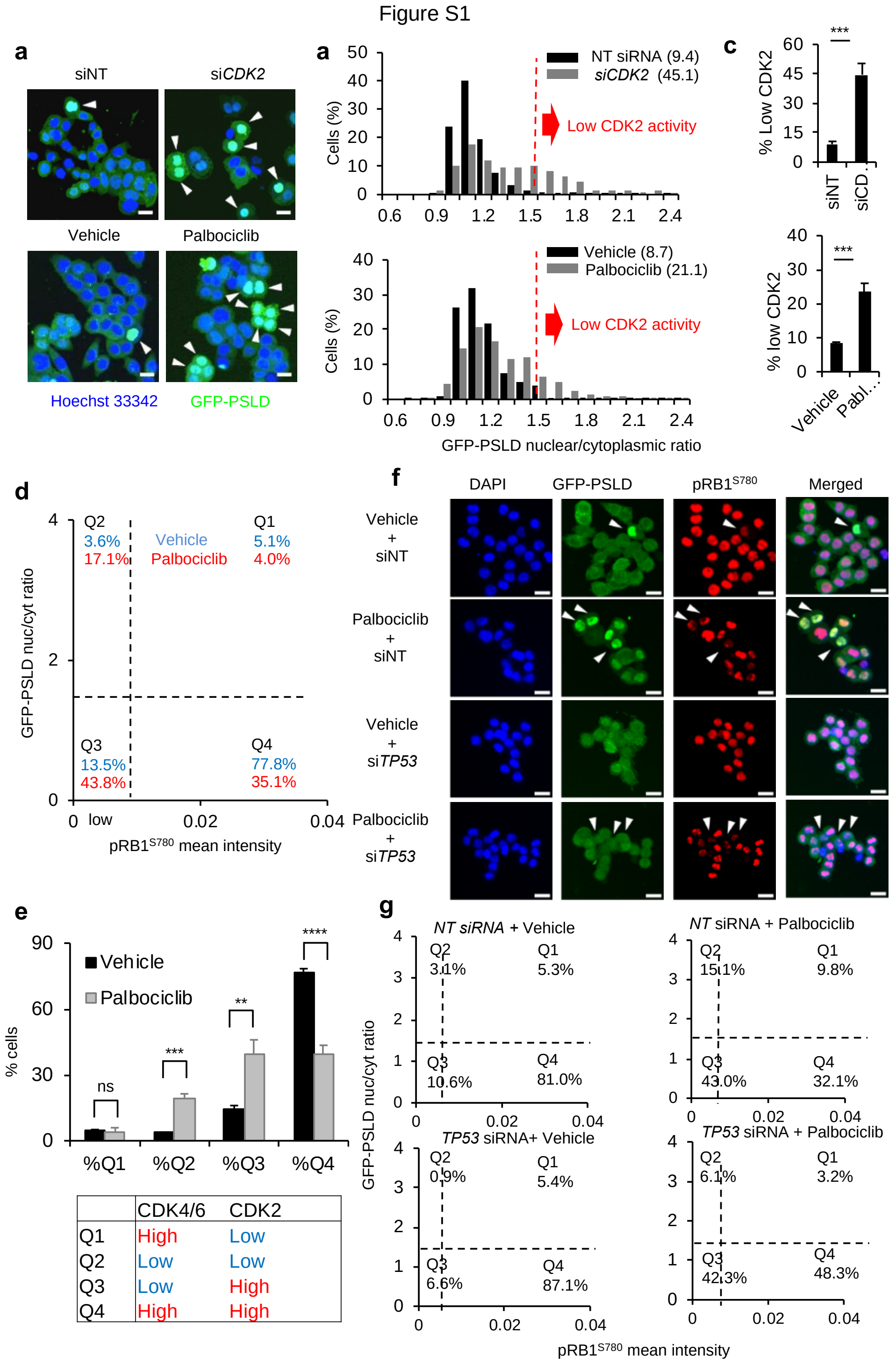


| | CDK4/6 | CDK2 |
|---|---|---|
| Q1 | High | Low |
| Q2 | Low | Low |
| Q3 | Low | High |
| Q4 | High | High |

**Figure S1.** Regulation of GFP-PSLD CDK2 reporter localization in HCT116.

**a** GFP-PSLD localisation in HCT116-PSLD transfected with *CDK2*-targeted siRNA (siCDK2) or non-targeted siRNA (siNT) or treated with 500 nM palbociclib or vehicle, for 48 h. Arrows mark cells with predominantly nuclear GFP-PSLD, indicative of low CDK2 activity.

**b** Population distribution of GFP-PSLD nuclear per cytoplasmic (nuc/cyto) fluorescence intensity ratios. Data were collected for > 2000 individual cells treated as in **a**. The dotted red lines identify the threshold for cells with low CDK2 activity (GFP-PSLD nuc/cyto ratio > 1.5). Numbers in parentheses show the percentage of cells in each treatment group for GFP-PSLD nuc/cyto ratio > 1.5**.**

**c** Graph depicting the fraction of cells with GFP-PSLD nuc/cyto ratio > 1.5 (low CDK2). Data represent mean ±SD for n = 3 independent repeats. Cells treated as in **a**; $^{**}p \leq 0.01$, $^{***}p \leq 0.001$, $^{ns}p > 0.05$, 2-tailed Student's t-test.

**d** Population scatter plot, depicting the relationship between CDK2 activity, assessed using the GFP-PSLD reporter, and CDK4/6 activity, assessed using immunostaining for $pRB1^{S780}$. Dotted lines represent thresholds used to identify cell populations with low CDK2 (GFP-PSLD nuc/cyto ratio > 1.5) and low $pRB1^{S780}$**.** Data represent **>** 2000 cells per condition.

**e** Graph displaying the fraction of cells within quadrants depicted in **d**. Data shown represent mean values ±SD, for n=3 independent repeats, $^{**}p \leq 0.01$, $^{***}p \leq 0.001$, $^{ns}p > 0.05$, $^{***}p \leq 0.001$, 2-tailed Student's t-test.

**f** Representative raw images of HCT116-PSLD, transfected with siRNA without target (siNT) or targeting *TP53* (siTP53) for 24 h followed by treatment with 500 nM palbociclib or vehicle for 24 h and stained with antibody for $pRB1^{S780}$. Arrows point to cells with low levels of $pRB1^{S780}$, indicative of loss of CDK4/6 activity.

**g** Population scatter plots comparing $pRB1^{S780}$ staining intensity and GFP-PSLD nuc/ cyto partitioning for > 2000 cells shown in **d.** Dotted lines represent thresholds that identify populations with low CDK2 (nuc/cyto ratio > 1.5) and low $pRB1^{Ser780}$.

Scale bar = 20 µm (**a**, **f**).

(Related to Figure 1)

Figure S2

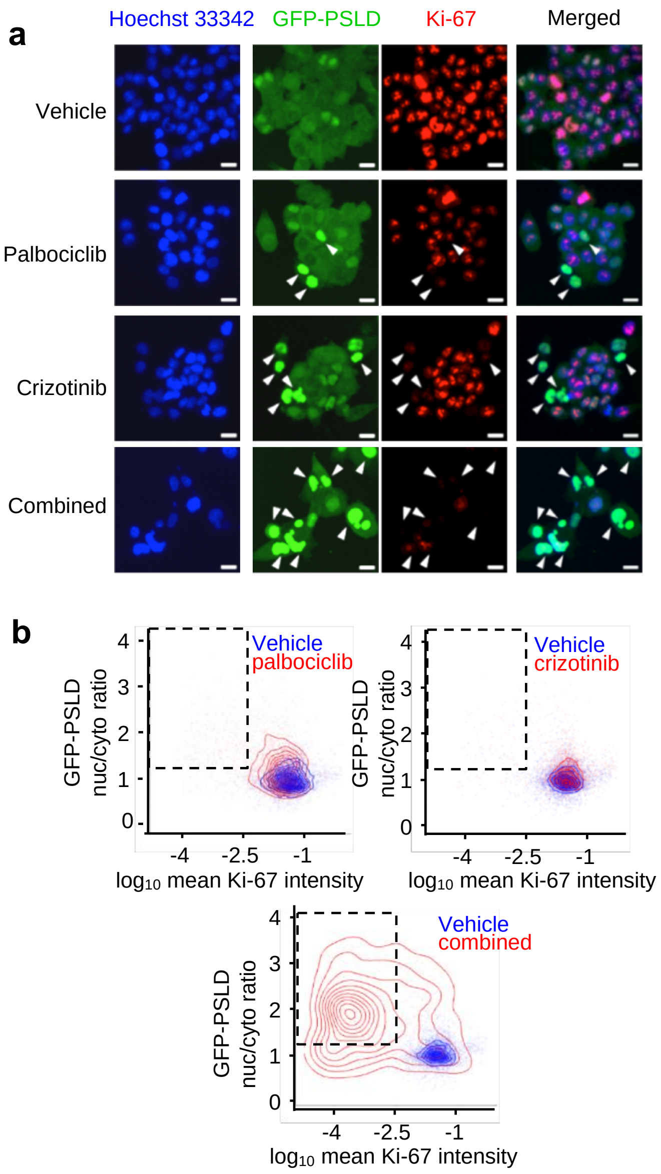


**c**

| | Combination | $CI_{50}$ | $CI_{75}$ | $CI_{90}$ |
|---|---|---|---|---|
| HCT116 $TP53^{WT}$ | Foretinib + Palbociclib | 0.19 | 0.12 | na |
| | Crizotinib + Abemaciclib | 0.49 | 0.47 | 0.49 |
| | Foretinib + Abemaciclib | 0.36 | 0.32 | 0.32 |
| HCT116 $TP53^{-/-}$ | Foretinib + Palbociclib | 0.12 | na | na |
| | Crizotinib + Abemaciclib | 0.04 | na | na |
| | Foretinib + Abemaciclib | 0.48 | 0.49 | na |

Colour scale for CI

0.01 — 1.0 — 2.0

Synergism — Additive — Antagonism

**Figure S2.** Inhibition of CDK4/6- and MET synergises to reduce the Ki-67 labeling index. **a** Representative raw images of HCT116-PSLD stained using anti-Ki-67 antibody following treatment with 450 nM inhibitors for 96 h. Arrows indicate cells with low CDK2 activity (GFP-

PSLD nuc/cyto ratio of > 1.5). Note low anti-Ki-67 fluorescence in cells with low CDK2 (GFP-PSLD nuc/cyto ratio of > 1.5).

**b** Population density plots depicting anti-Ki-67 signal intensity and GFP-PSLD localization in cells populations treated as in **a**. Dotted boxes identify the cell population with low CDK2 activity (GFP-PSLD nuc/cyto ratio of > 1.5) combined with low anti-Ki-67 signal intensity.

**c** Synergistic reduction of the Ki-67 labelling index by chemically distinct CDK4/6 and MET inhibitors. Cells were treated with inhibitors for 96 h. Results for $TP53^{WT}$ (upper) and $TP53^{-/-}$ (lower) HCT116 are shown. Data represent mean of 2 independent repeats, run in duplicate each. $CI_{50}$, $CI_{75}$ and $CI_{90}$ denote CI values at concentrations yielding a reduction in Ki-67 labelling index by 50%, 75% or 90%, respectively, na= fractional responses not achievable within the inhibitor concentration range tested.

(Related to Figure 3)

Figure S3

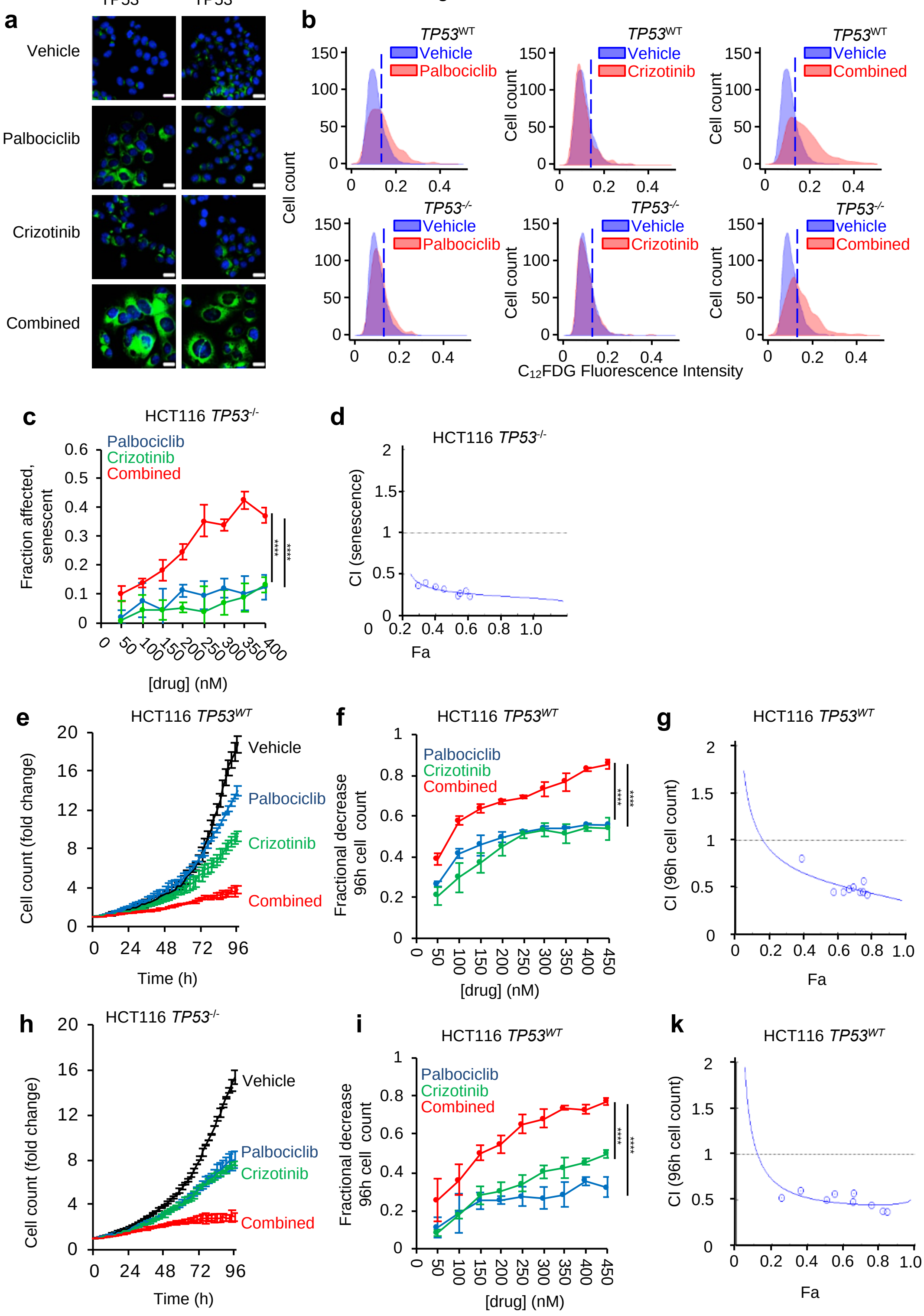


**Figure S3.** CDK4/6 and MET inhibitors cooperate to increase the fraction of senescence-marker positive cells and to decrease population growth rates.

**a** Effect of combined inhibitor treatment on the expression of senescence marker SA-β-gal**.** Representative images of HCT116 treated with 500 nM inhibitors for 120 h and detection of SA-β-gal activity by incubation with $C_{12}$FDG at acidic pH. Scale bar = 20 µm.

**b** Distribution of perinuclear $C_{12}$FDG- fluorescence intensity in cells treated as in **a**. Dotted lines indicated threshold used to identify cells with high SA-β-gal activity. Results for $TP53^{WT}$ and $TP53^{-/-}$ HCT116 are shown.

**c and d** Concentration-effect analysis depicting the fractional increase in cells with high SA-β-gal activity for HCT116 $TP53^{-/-}$. Fractional increase in cells with high SA-β-gal (**c**) and CI value plots (**d**). Data represent means ±SD for n= 3 independent repeats, run in tripicate each. ****$p \leq 0.0001$, 2-way ANOVA comparing the effects size of single agent against that of the combination.

**e-k** Effect of inhibitor treatment on cell population growth determined using quantitative time-lapse microscopy**.** (**e** and **h**) Graphs depicting trace of cell number over time using 250 nM of each inhibitor. (**f** and **i**) Concentration-effect analysis depicting the fractional decrease in cell numbers relative to vehicle at 96 h. Data are means±variance for 2 independent repeats, with triplicates each. (**g**, **k**) CI value plots relating to **f** and **i**. ****$p \leq 0.0001$, 2-way ANOVA comparing omparing the effects size of single agent against that of the combination.

(Related to Figure 4)

Figure S4

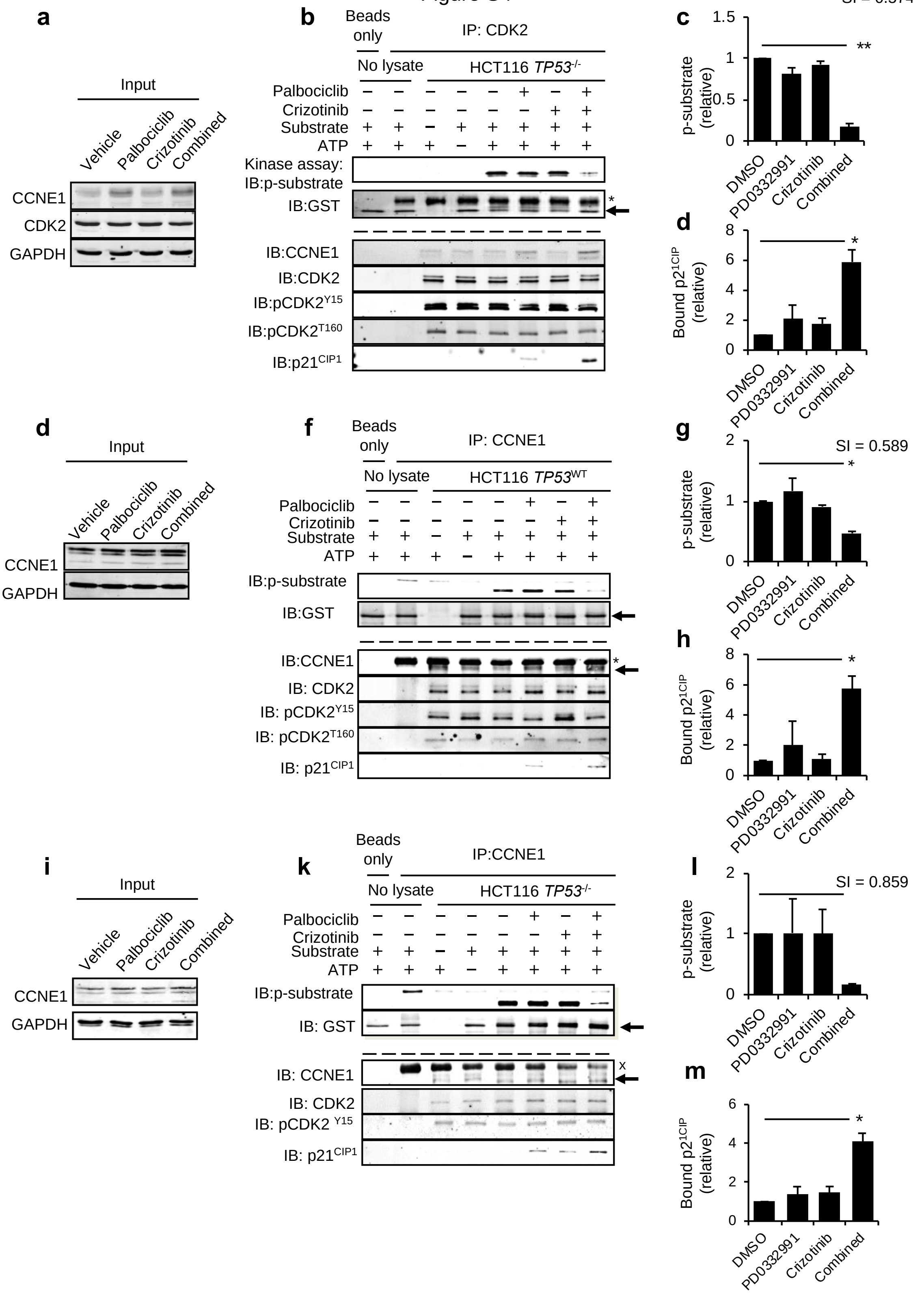


**Figure S4.** Co-operative control of CDK2 by MET and CDK4/6 inhibitors involves p21$^{CIP1}$. Characterisation of CDK2 or CCNE1 complexes in inhibitor treated cells using immunoprecipitation. Lysates were generated after cells were treated with 500 nM of inhibitors for 24 h.

**a–d** Data for anti-CDK2 precipitates. Data for HCT116 *TP53*$^{-/-}$ cells treated with 500 nM inhibitor for 24 h (**a**) Immunoblot of input lysates. (**b**) Immunoprecipitation kinase assay, abundance of *in vitro* phosphorylated GST-pRB 763-928 substrate (p-substrate) reflecting CDK2 activity and total substrate (upper), and abundance of p21$^{CIP1}$, total and phosphorylated CDK2 (pCDK2$^{Y15}$, pCDK2$^{T160}$), and CCNE1 in the respective immunoprecipitations, as indicated (lower). (**c**) Mean abundance (± range) of p-substrate and (**d**) mean abundance of co-precipitated p21$^{CIP1}$ relative to vehicle-treated cells. Data represent mean of 2 independent repeats. SI values were calculated using mean values. Precipitation reaction in the absence of antibody (Bead only) or absence of lysate (No lysate) as indicated. *$p \leq 0.05$, ** $p \leq 0.01$, 2-way ANOVA assessing the effect size of single agent with effect size of their combination.

**e–m** Data using anti-CCNE1 precipitation, using HCT116 *TP53*$^{W}$ (**e-h**) or HCT116 *TP53*$^{-/-}$ cells (**i-m**). Experimental design and analysis was as for **a-d.** Arrows, where used, denote the position of antibody detected proteins, $^{X}$ denotes the position of IgG. Data (**g** and **h**, **l** and **m**) represent n = 2 independent repeats. *$p \leq 0.05$, 1-way ANOVA.

(Related to Figure 5)

Figure S5

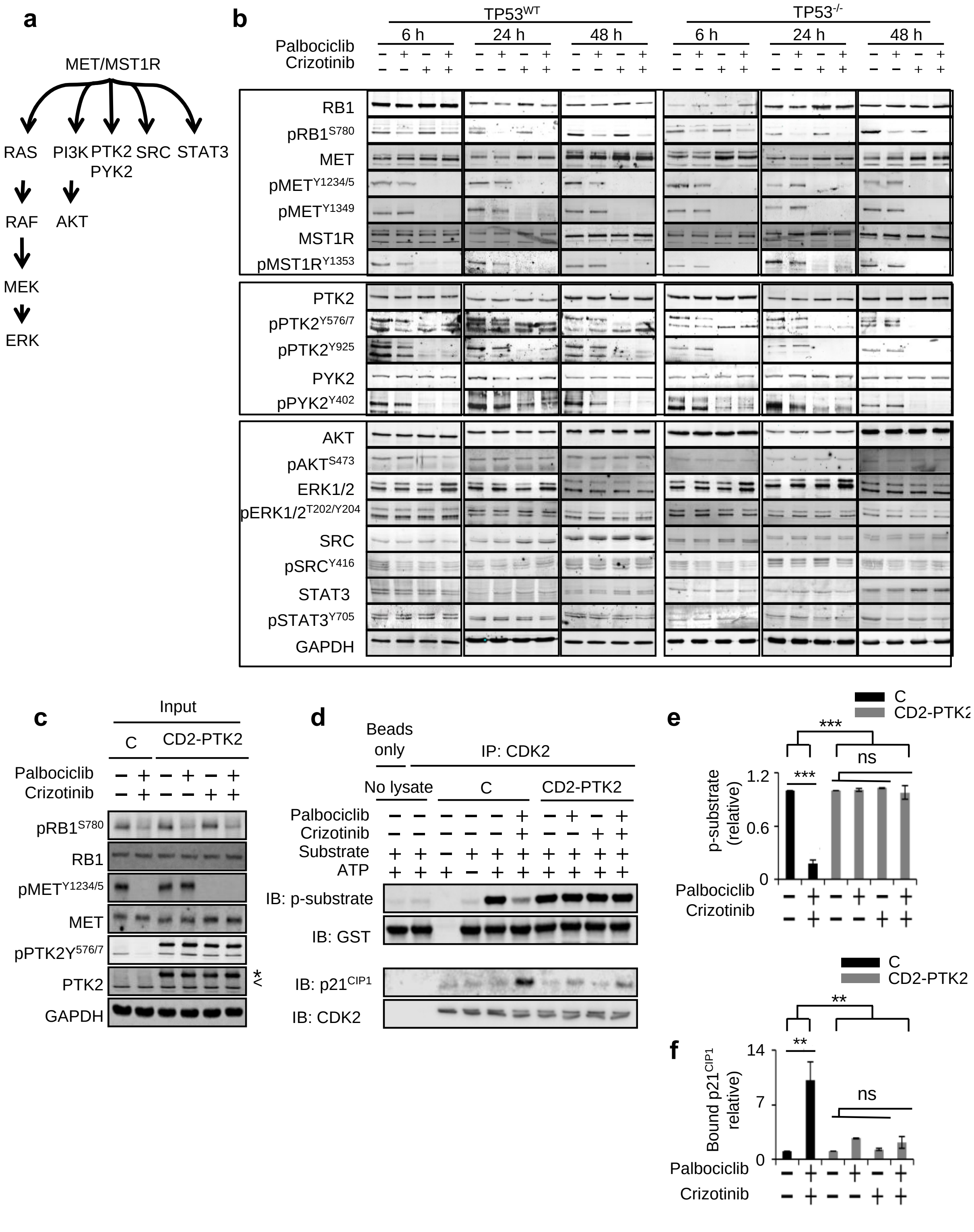


**Figure S5:** Role of MET effector FAK family kinases in permitting CDK4/6-independent CDK2 activation.

**a** Schematic depicting effector pathways for MET and MST1R.

**b** Abundance and activation state of MET effectors in lysates of cells treated with 500 nM inhibitors. Shown are total and phosphorylated RB1 and MET family proteins (upper panel), total and phosphorylated FAK family kinases (middle panel) and total and phosphorylated forms of MET effectors AKT, ERK1/2, SRC and STAT3 (lower panel).

**c–e** Characterisation of CDK2 complexes in HCT116 *TP53*$^{-/-}$ cells expressing constitutively active PTK2 (CD2-PTK2). Cells were treated with 500 nM inhibitors for 24 h. (**c**) Immunoblot of input lysates. < identifies signal for cell-encoded PTK2, * identifies CD2-PTK2. (**d**) Anti-CDK2 immunoprecipitation kinase assays: (upper) abundance of *in vitro* phosphorylated substrate GST-pRB 763-928 (p-substrate), and total substrate, (lower) abundance of p21$^{CIP1}$ and CDK2 in the same immunoprecipitation reaction. (**e**) Mean abundance of p-substrate and (**f**) mean abundance of co-precipitated p21$^{CIP1}$ relative to vehicle-treated cells, for 3 independent repeats. ***$p \leq 0.01$, ** $p \leq 0.001$ 1-way ANOVA comparing value for vehicle and the combination in control cells. $^{ns}p > 0.05$, 2–way ANOVA assessing effect size of single agent compared with effect size of the combination in HCT116 CD2-PTK2, and ***$p \leq 0.01$, ** $p \leq$ 0.001, 2-way ANOVA assessing the effect size of the combination in HCT116 control and HCT116 CD2-PTK2. C = control cells (**c–f**).

(Related to Figure 6)

Figure S6

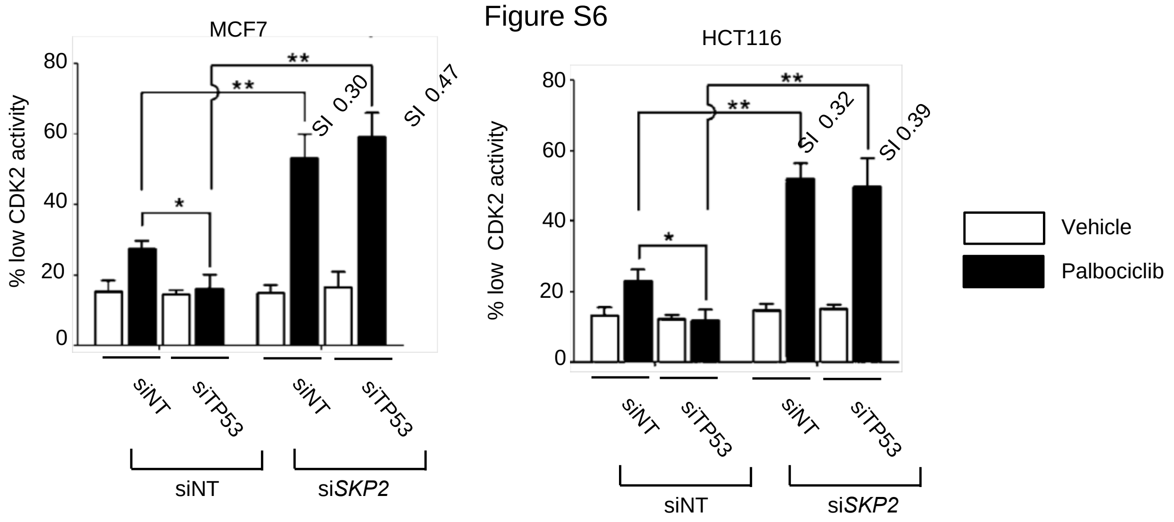


**Figure S6.** SKP2 Loss co-operates with palbociclib to decrease CDK2 activity. Charts show the percentage of cells with a GFP-PSLD nuc/cyto ratio of > 1.5, indicative of low CDK2 activity. Cells were transfected with siRNA combinations as indicated 24 h followed by treatment with 500 nM palbociclib or vehicle for a further 24 h. Results for HCT116 and MCF7 cells are shown. Data are mean values ±SD for n= 3 independent repeats. *$p \leq 0.05$, **$p \leq 0.01$, 2-way ANOVA coparing the effect size in TP53 compromised versuys T53 normal cells and comparing the effect size of palbociclib vs vehicle in unperturbed (siNT) and SKP2 ablated (siSKP2) cells, SI values for palbociclib + si*SKP2* are indicated.

### Supplementary tables

**Supplemental Table S1:** Results of MetaCore™ pathway map enrichment analysis using hits identified in *TP53*[KD] cells.

| Rank | Name of MetaCore™ pathway map | No. of kinases in pathway | No. of hits | p-value | Hits mapped |
|---|---|---|---|---|---|
| 1 | MET and MSP receptor (RON) signalling pathways in SCLC | 18 | 5 | $3.00\times10^{-5}$ | *MST1R, PTK2B, MET, CDK2, PTK2* |
| 2 | SDF-1 signalling in hematopoietic stem cell homing | 11 | 3 | $1.73\times10^{-3}$ | *PTK2B, PTK2, JAK2* |
| 3 | The role of PTEN and PI3K signalling in melanoma | 13 | 3 | $2.92\times10^{-3}$ | *BRAF, PTK2, JAK2* |
| 4 | HBV-dependent NF-kB and PI3K/AKT pathways leading to HCC | 13 | 3 | $2.92\times10^{-3}$ | *PTK2B, CDK2, PTK2* |
| 5 | FGFR3 signalling in multiple myeloma | 14 | 3 | $3.66\times10^{-3}$ | *PTK2B, FGFR3, JAK2* |
| 6 | Resolution of inflammation in healing myocardial infarction | 5 | 2 | $5.34\times10^{-3}$ | *MET, JAK2* |
| 7 | Proliferative action of gastrin in pancreatic cancer | 16 | 3 | $5.44\times10^{-3}$ | *PTK2B, PTK2, JAK2* |
| 8 | CXCR4 signalling pathway | 16 | 3 | $5.44\times10^{-3}$ | *PTK2B, PTK2, JAK2* |
| 9 | Growth hormone signalling via PI3K/AKT and MAPK cascades | 17 | 3 | $6.54\times10^{-3}$ | *PTK2B, PTK2, JAK2* |
| 10 | Cadherin-mediated cell adhesion | 6 | 2 | $7.91\times10^{-3}$ | *MET, PTK2* |

**Supplemental Table S2:** Results of MetaCore™ pathway map enrichment analysis using hits identified in *TP53*[WT] cells.

| Rank | Name of MetaCore™ pathway map | Total kinases in pathway | No. of hits | p-value | Hits mapped |
|---|---|---|---|---|---|
| 1 | Role of SCF complex in cell cycle regulation | 9 | 4 | $7.53\times10^{-5}$ | *PLK1, CDK2, CKS1B, CDK1* |
| 2 | Role of APC in cell cycle regulation | 9 | 4 | $7.53\times10^{-5}$ | *PLK1, CDK2, CKS1B, CDK1* |
| 3 | Cell cycle progression in prostate cancer | 11 | 4 | $1.89\times10^{-4}$ | *CDK2, RPS6KB1, CDK1, JAK2* |
| 4 | nNOS signalling in neuronal synapses | 2 | 2 | $9.08\times10^{-4}$ | *DLG4, CALM3* |
| 5 | Abnormalities in cell cycle in SCLC | 9 | 3 | $1.88\times10^{-3}$ | *CDK2, CKS1B, CDK1* |
| 6 | Leptin signalling in colorectal cancer | 10 | 3 | $2.63\times10^{-3}$ | *RPS6KB1, CDK1, JAK2* |
| 7 | Constitutive and activity-dependent synaptic AMPA receptor delivery | 10 | 3 | $2.63\times10^{-3}$ | *DLG4, DLG1, CALM3* |
| 8 | Transition and termination of DNA replication | 3 | 2 | $2.67\times10^{-3}$ | *CDK2, CDK1* |
| 9 | Main chemotherapy drugs and their action in SCLC cells | 24 | 4 | $4.71\times10^{-3}$ | *ABL1, CDK2, RPS6KB1, CDK1* |
| 10 | Role of nicotine-induced leptin resistance in hypothalamus in development of obesity | 4 | 2 | $5.25\times10^{-3}$ | *RPS6KB1, JAK2* |
